# Defining the ultrastructure of the hematopoietic stem cell niche by correlative light and electron microscopy

**DOI:** 10.1101/2020.11.12.380790

**Authors:** Sobhika Agarwala, Keun-Young Kim, Sebastien Phan, Saeyeon Ju, Ye Eun Kong, Guillaume A. Castillon, Eric A. Bushong, Mark H. Ellisman, Owen J. Tamplin

## Abstract

The blood system is supported by hematopoietic stem and progenitor cells (HSPCs) found in a specialized microenvironment called the niche. Many different niche cell types support HSPCs, however how they interact and their ultrastructure has been difficult to define. Here we show that single endogenous HSPCs can be tracked by light microscopy, then identified by serial block-face scanning electron microscopy (SBEM) at multiscale levels. Using the zebrafish larval kidney marrow (KM) niche as a model, we followed single fluorescently-labeled HSPCs by light sheet microscopy, then confirmed their exact location in a 3D SBEM dataset. Our approach allowed us to identify dopamine beta-hydroxylase (dbh) positive ganglia cells as a previously uncharacterized functional cell type in the HSPC niche. By integrating multiple imaging modalities, we could resolve the ultrastructure of single rare cells deep in live tissue and define all contacts between an HSPC and its surrounding niche cell types.

## Introduction

Hematopoietic stem and progenitor cells (HSPCs) give rise to all blood cell types throughout the life of an organism (Orkin and Zon, 2008). HSPCs reside in a complex microenvironment called the niche that is made up of many different kinds of support cells, including various types of mesenchymal stromal cells (MSCs) and endothelial cells (ECs) (Pinho and Frenette, 2019). However, our understanding of HSPC interactions with niche cells has been limited to low resolution light microscopy. Recently developed transgenic reporter lines have allowed identification of well-defined endogenous HSPCs in both mouse and zebrafish model organisms (Acar et al., 2015; Chen et al., 2016; Christodoulou et al., 2020; Tamplin et al., 2015). The dynamic behavior of HSPCs in the niche of mouse and zebrafish can be observed using live imaging (Bixel et al., 2017; Christodoulou et al., 2020; Itkin et al., 2016; Koechlein et al., 2016; Lo Celso et al., 2009; Spencer et al., 2014; Tamplin et al., 2015). Yet it remains challenging to observe HSPC-niche interactions at high resolution. Certain events during hematopoietic ontogeny, such as the colonization of the fetal bone marrow, have so far only been studied in fixed tissues because they are difficult to access (Coskun et al., 2014). Our goal is to better define the ultrastructure of single endogenous HSPCs deep in niche tissue.

We have taken advantage of the transparency and external development of the zebrafish larva to study the earliest migration events of HSPCs into the presumptive adult kidney marrow (KM) niche. We considered using a correlative light and electron microscopy (CLEM) (Karreman et al., 2016) to resolve the ultrastructure of single lodged HSPCs. To confirm cell identity we used a label-based approach similar to what was done using an APEX2-Venus-CAAX fusion protein (Hirabayashi et al., 2018) that could be visualized both with light microscopy and high-contrast in EM imaging. A similar method was applied in zebrafish using an APEX-GBP (GFP Binding Protein) fusion to resolve GFP^+^ transgene expression on electron micrographs (Ariotti et al., 2015). Building on these previous studies, we used genetically encoded APEX2 engineered peroxidase (Lam et al., 2015), together with a fluorescent protein, to track single HSPCs as they migrated into and lodged in the larval KM during the earliest colonization stages. We found clusters of HSPCs around the glomerulus as previously described (Murayama et al., 2006), as well as single HSPCs lodged in a perivascular niche. Using 3D datasets generated by SBEM, we could confirm that single HSPCs were directly and simultaneously contacted by multiple niche cell types, including MSCs, ECs, and a previously uncharacterized ganglia-like cell in the niche.

## Results

To determine the location and quantify all HSPCs in the KM, we imaged fixed, optically cleared HSPC-specific Runx:mCherry^+^ transgenic zebrafish (Tamplin et al., 2015) larvae at 5 days post fertilization (d.p.f.). HSPCs were seen in bilateral clusters of ~50 cells on either side of the glomerulus, mediolateral to the proximal pronephric tubules (Extended Data Figure 1A-D). In addition, the HSPCs were not highly proliferative as seen by phospho-histone H3 (PH3) antibody labeling (Extended Data Figure 1E-I), suggesting that cells in the larval KM niche are relatively quiescent at 5 d.p.f.; similar results were previously observed at 7.5 d.p.f. (van Rooijen et al., 2009).

To follow the dynamics of HSPC colonization in the KM niche, we performed time-lapse live imaging. Although the early zebrafish larva is transparent, imaging the KM using point scanning confocal microscopy is challenging due to the relatively slow acquisition time and depth of the tissue. To rapidly capture HSPC colonization events throughout the entire larval KM, we performed light sheet fluorescence microscopy (Huisken and Stainier, 2009). Using this technique, we could rapidly acquire a Z-stack through the entire KM in less than 30 seconds (>200 slices with 1 μm spacing). Time-lapse images showed Runx:mCherry^+^ HSPC clusters mediolateral to the proximal pronephric tubules (Supplementary Video 1), confirming our observations in fixed embryos. Together with a reporter line for pronephric tubules (cdh17:GFP (Zhou et al., 2010); Extended Data Figure 2), we localized Runx:mCherry^+^ HSPCs within the anterior kidney region. To determine the location of these HSPC clusters relative to the vasculature, we similarly imaged HSPC-specific Runx:mCherry together with the flk:ZsGreen vascular-specific transgenic reporter (Cross et al., 2003). HSPC clusters were located between lateral dorsal aortae and cardinal veins (Extended Data Figure 3). Light sheet live imaging of the KM niche at 5 d.p.f. allowed us to track the very early dynamics of HSPC colonization.

To more specifically observe the interaction between single HSPCs and niche, we performed similar imaging using the Runx:GFP transgenic line together with flk:mCherry to label vessels (Supplementary Video 2). Runx:GFP is a more restricted marker of HSPCs compared to Runx:mCherry, which more broadly labels the progenitor pool (Tamplin et al., 2015) (Extended Data Figure 4). Upon arrival within the KM, rare circulating GFP^+^ HSPCs were seen interacting with and lodging in the posterior perivascular niche (Figure 1A). We could resolve that lodged cells were surrounded by ECs in a pocket-like structure, as we observed previously in the zebrafish caudal hematopoietic tissue (CHT) and mouse fetal liver (Tamplin et al., 2015) (Figure 1B).

**Figure 1.**
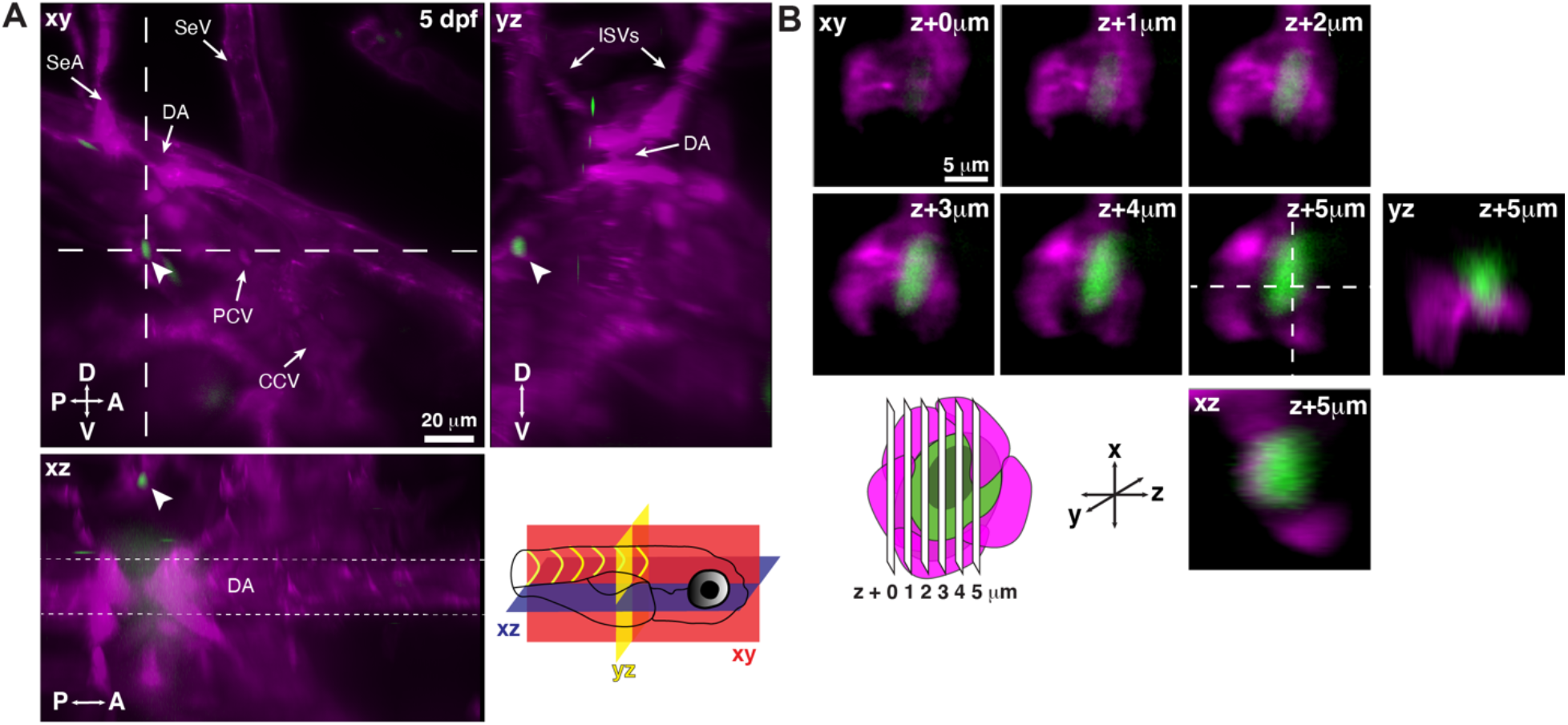
Single HSPCs lodge in a perivascular region of the larval kidney niche. **(A)** Snapshot of single optical sections (xy, xz, yz planes) from light sheet live image of a Runx:GFP;flk:mCherry double transgenic zebrafish larva. A single Runx:GFP^+^ HSPC (white arrowhead) is lodged in the perivascular region lateral to the dorsal aorta (DA). **(B)** Detail of optical sections (1 μm steps) through the single lodged Runx:GFP^+^ HSPC in (A). mCherry^+^ endothelial cells contact the HSPC and form a surrounding pocket. The +5 μm section is also shown in xz and yz planes. Abbreviations: DA, dorsal aorta; SeA, intersegmental artery; SeV, intersegmental vein; PCV, posterior cardinal vein; CCV, common cardinal vein; ISVs, intersegmental vessels; D, dorsal; V, ventral; A, anterior; P, posterior.

We investigated whether remodeling of niche cells around HSPCs leads to localization of tight junction scaffolding protein such as Zonula occludens-1 (ZO-1) (Anderson et al., 1988; Stevenson et al., 1986) at the cell junctions. In sections obtained from Oregon green injected Runx:mCherry^+^ transgenic larvae, we observed that the regions surrounding the HSPCs were enriched for ZO-1 antibody (Figure 2A,B). This suggests that ZO-1 is a potential candidate for mediating adhesion between HSPCs and the surrounding niche cells in the KM niche.

**Figure 2.**
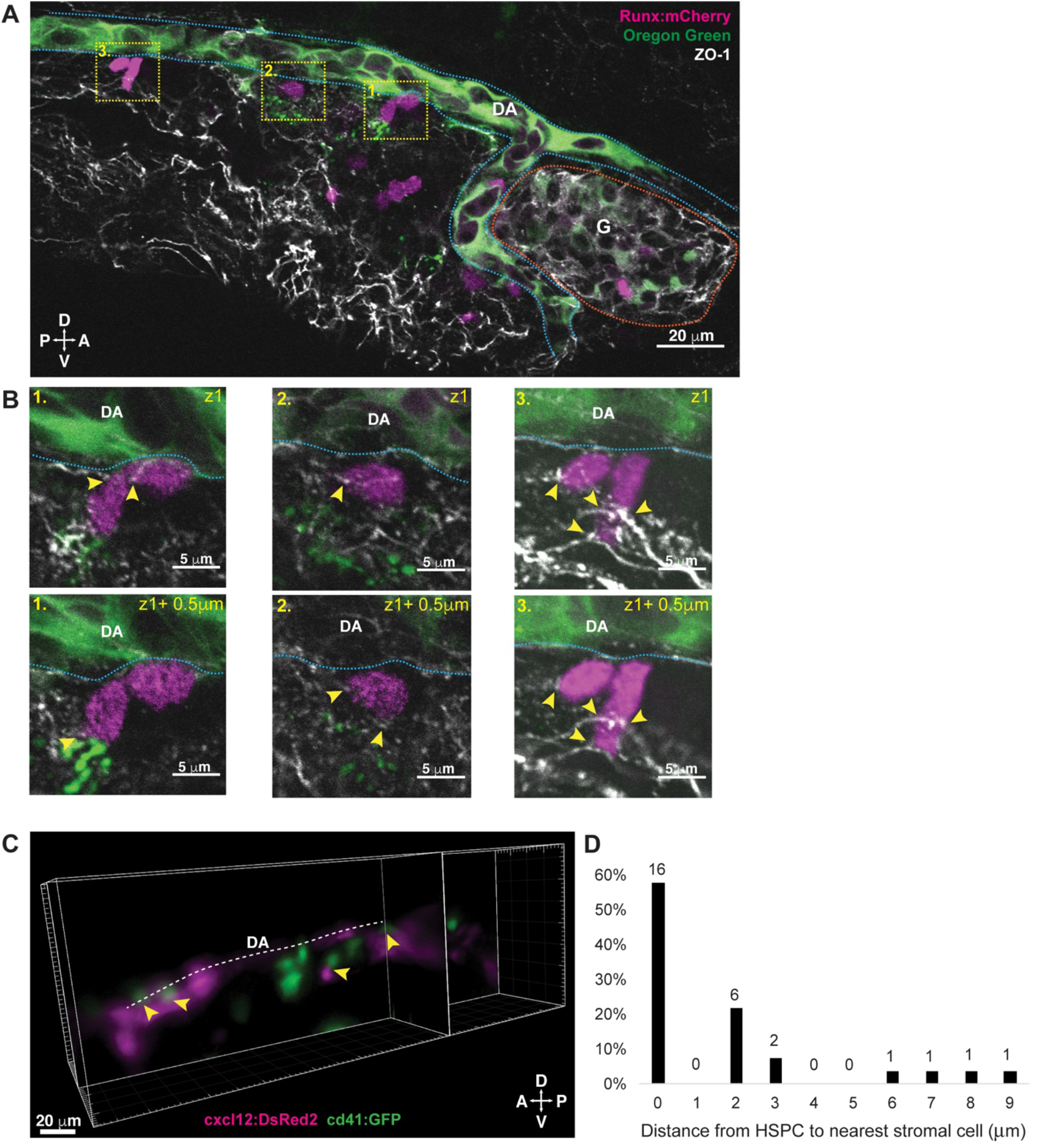
HSPCs lodged in the larval kidney niche make direct contacts with ECs and MSCs. **(A)** Single optical section from confocal image of larval kidney (fixed) shows Runx:mCherry^+^ HSPCs (magenta) lodged in the perivascular niche. Oregon Green dye labels the vessel lumen. Blue dotted lines surround the dorsal aorta (DA) and red dotted lines surround the glomerulus (G). Tight junction protein is marked by ZO-1 (white). **(B)** High resolution optical sections (0.5 μm steps) through the boxed regions in (A) show ZO-1^+^ contact points between mCherry^+^ HSPCs and the niche (yellow arrowheads). **(C)** Orthogonal slices (xy and yz planes) from live light sheet 3D volume of larval kidney niche. Single cd41:GFP^+^ HSPCs (green) are in contact or in close proximity (yellow arrowheads) to cxcl12:DsRed2^+^ MSCs (magenta). Dotted line represents the dorsal aorta (DA). **(D)** Quantification of distances measured between GFP^+^ HSPC and DsRed2^+^ MSCs shows ~60% of HSPCs are in contact with MSCs, and the remaining are within 9um. Numbers above the columns indicate the cell numbers counted in each group (from n = 8 embryos). D, dorsal; V, ventral; A, anterior; P, posterior.

We performed imaging using the cd41:GFP hematopoietic transgenic line together with cxcl12:DsRed2 to label MSCs (Glass et al., 2011) (Figure 2C). We measured the distance between HSPCs and MSCs and observed 60% of HSPCs were in direct contact with MSCs, and all other HSPCs were within one cell diameter (~9 μm) of an MSC (Figure 2D). Together, our results demonstrate that HSPC lodgement in the larval KM niche occurs close to or in contact with a single MSC, and that ECs surround the HSPC, similar to what we observed previously in the CHT (Tamplin et al., 2015).

To understand the endogenous microenvironment of this perivascular region of the KM, we needed to develop an approach to resolve the ultrastructure of a single lodged HSPC and its surrounding niche cells. Zebrafish larvae have been analyzed using a serial section electron microscopy (EM) technique that resolved the projectome of the complete brain (Hildebrand et al., 2017). We previously used CLEM based entirely on anatomical landmarks to match the position of fluorescently labeled HSPCs in the CHT niche between confocal and SBEM datasets (Tamplin et al., 2015). However, we found this approach was difficult to apply in the KM because the tissue is much larger and denser than the CHT (data not shown). To fully validate correlation of single cells between different imaging modalities (i.e. light and electron microscopy), we used genetically encoded APEX2 to label single endogenous HSPCs. In this way we were able to track HSPC lodgement in the perivascular KM niche, then resolve 3D ultrastructure by SBEM at nanoscale.

To achieve these experimental goals, we developed the following workflow. First, we generated a transgenic construct that expressed mCherry for light microscopy, and APEX2 (enhanced ascorbate peroxidase) as a genetic tag that allow electron-dense contrast on target subcellular structure (Figure 3A; APEX2-H2B and mito-APEX2 for localization to the nucleus and the mitochondrial matrix, respectively). To drive expression of this construct in HSPCs, we cloned these regions under control of the *draculin* promoter, a marker for vascular and hematopoietic lineages (Mosimann et al., 2015). The rationale for our approach was that in transgenic larvae, HSPCs that are mCherry^+^ will also be APEX2^+^, making them identifiable in both fluorescence and EM, respectively (Figure 3A). To track and correlate these single cells through multiple imaging modalities, we required sparse labelling of HSPCs. Therefore, we generated transient F0 transgenics with a mosaic-labeled HSPC pool. We first calculated the proportion of mCherry^+^;APEX2^+^ cells that co-localized with cd41:GFP^+^ cells by injecting the *draculin:mito-APEX2_p2A_APEX2-H2B_p2A_mCherry* (referred hereafter as *draculin:2APEX2_mCherry*^+^) construct together with *tol2* transposase into the *cd41:GFP* transgenic line. We observed that 15% of HSPCs were both mCherry^+^;APEX2^+^ and GFP^+^ (data not shown). Specifically, in draculin:2APEX2_mCherry^+^ transients alone, we detected 1-2 mCherry^+^ cells/EC pockets and 1-5 mCherry^+^/clusters in 11 larvae. This degree of labeling was similar to that of a stable transgenic HSPC line like cd41:GFP^+^ where we detected 1-2 GFP^+^ cells/EC pockets and 4-9 GFP^+^ cells/clusters in 6 larvae (data not shown). In the Runx:GFP^+^ HSPC reporter line there were 1-2 GFP^+^ cells/EC pockets in 8 larvae (Figure 1). This suggests that transient draculin expression generates sparsely labelled HSPCs that lodge within the niche pocket with the same frequency as seen for other stable hematopoietic lines.

**Figure 3.**
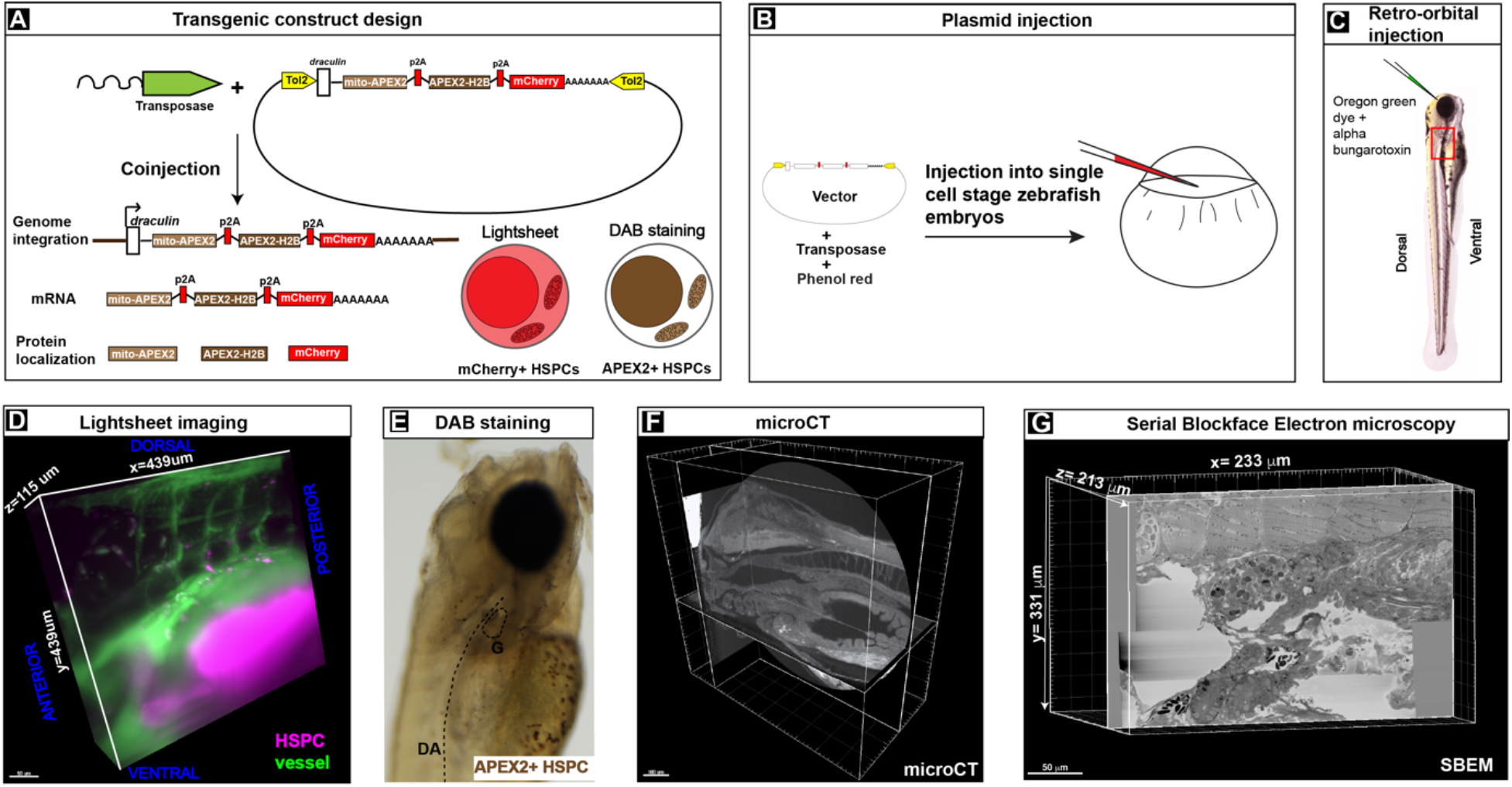
Workflow to genetically encode a label in endogenous HSPCs for live tracking by LM and high contrast resolution in SBEM sections. **(A)** Fusion construct encoding p2A-linked proteins mito-APEX2, APEX2-H2B, and mCherry that localize to the mitochondria, nucleus, and cytoplasm, respectively. The *draculin* promoter was used to transiently drive strong mosaic expression in HSPCs. Random insertion in the genome was by Tol2-mediated transgenesis. **(B)** Tol2 *draculin:mito-APEX2_p2A_APEX2-H2B_p2A_mCherry* fusion construct was injected together with *tol2* mRNA in one cell wild type zebrafish embryos. **(C)** At 5 d.p.f., embryos with circulating mCherry^+^ HSPCs were visually screened and retro-orbitally injected with alpha bungarotoxin to paralyze the embryo, and Oregon Green dye to label the vasculature. **(D)** Dye-injected mCherry^+^ double positive embryos were visually screened and used for light sheet microscopy (example shows a 439×439×115 μm^3^ volume of the anterior KM). **(E)** Brightfield example of a single embryo after fixation and DAB staining to label APEX2^+^ HSPCs (G: glomerulus, DA: dorsal aorta). **(F)** After embedding, the sample was oriented and trimmed using microCT (example shows orthogonal sections in 3 planes). **(G)** Single plane from ~3,000 sections of SBEM data (example shows a 233×331×213 μm^3^ volume).

Once the identity of these mCherry^+^;APEX2^+^ cells were confirmed, this construct was co-injected together with *tol2* transposase into single-cell stage wild type zebrafish embryos (Figure 3B). At 5 d.p.f., larvae with circulating mCherry^+^ HSPCs were injected retro-orbitally with alpha-bungarotoxin and dextran-conjugated Oregon Green dye, to paralyze the larvae and label the vasculature, respectively (Figure 3C). Once immobilized and mounted for light sheet live imaging, optical sections were acquired through the entire depth of the KM. A short timelapse was performed to confirm an HSPC was lodged in the KM and not circulating (~30 minutes; Figure 4A; Supplementary Video 3). Single lodged HSPCs could be identified relative to the surrounding tissues (Figure 3D), immediately after imaging, larvae were fixed, and after DAB (3,3’-Diaminobenzidine) labeling reaction, APEX2^+^ cells were identified by brightfield microscopy (Figure 3E). Larvae were treated with osmium tetroxide and embedded for microcomputed tomography (microCT; Figure 3F and Extended data Figure 5; Supplementary Video 4). This intermediate microCT step allowed the larvae to be oriented and trimmed to select a discrete region of interest (ROI) for SBEM (Figure 3G and Extended data Figure 5). Lastly, automated SBEM generated over 3,000 high-resolution sections of the ROI (e.g. XY=10.8 nm/pixel, Z=70 nm/pixel; Extended data Figure 5; Supplementary Video 5). Using focal charge compensation (Focal CC) (Deerinck et al., 2018) which effectively eliminates specimen charging, we obtained a high-resolution SBEM dataset without the need for excessive postprocessing alignment. Together, these experimental steps allowed us to label an endogenous HSPC that was tracked live, then stained for high contrast detection in a large SBEM dataset.

**Figure 4.**
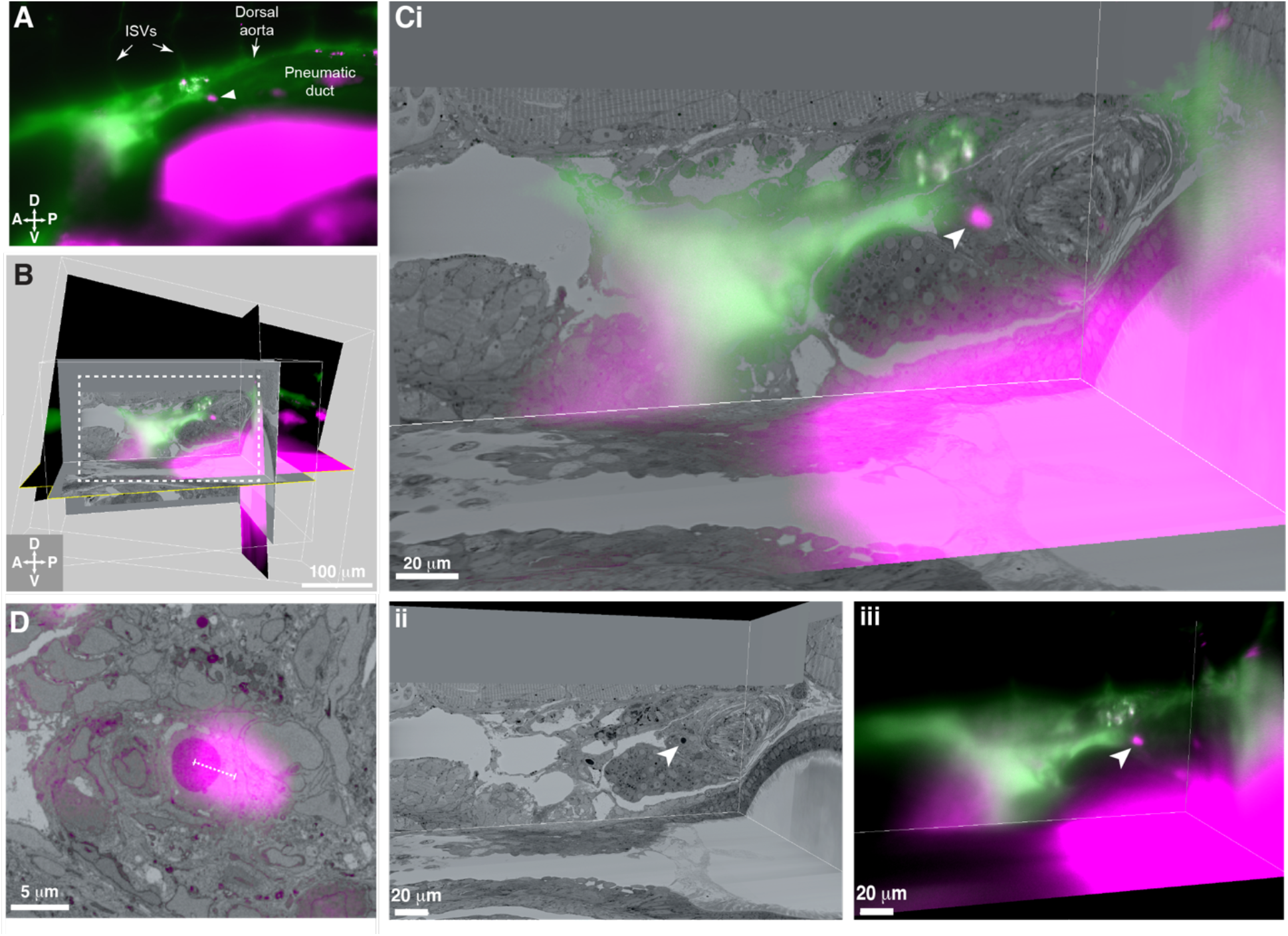
3D alignment of light sheet and SBEM datasets localizes a single rare HSPC across multiple imaging modalities. **(A)** Single z-plane from lightsheet imaging of *draculin:2APEX2_mCherry^+^* transgenic larva showing the lodged mCherry+ HSPC (white arrowhead; ISVs, intersegmental vessels). **(B)** Global alignment of 3D rendered models generated from light sheet and SBEM datasets using Imaris software. **(C)** Orthogonal views of the white boxed region within A shows a 3D view of the alignment between light sheet and SBEM datasets. (i) White arrowhead points to the single lodged HSPC in the aligned light sheet and SBEM datasets. (ii) APEX2^+^ HSPC in SBEM data. (iii) mCherry^+^ HSPC in light sheet data (iii). Green: Injected Oregon Green dextran dye marking vessels. Magenta: Runx:mCherry^+^ HSPCs and autofluorescence in gut. **(D)** Detail of the alignment shows mCherry^+^ HSPC and APEX2^+^ HSPC are <5 μm apart (dotted white line).

To correlate the position of a single labeled HSPC across multiple imaging modalities, we performed 3D software alignment of both light sheet and SBEM datasets. First, we identified anatomical features as landmarks in the SBEM data, such as the somites, glomerulus, and pronephric tubules (Extended data Figure 6). We also observed clustered hematopoietic cells around the glomerulus in the same region as seen by light sheet microscopy (Figure 1; Extended data Figure 7). We merged 3D rendered light sheet and SBEM datasets using image analysis software (Imaris) and aligned matching anatomical features in all three planes (Figure 4B,C; Extended data Figure 8). By performing these 3D alignments, we could locate a single APEX2^+^ cell in the SBEM dataset that was <5 μm from the corresponding mCherry^+^ HSPC imaged in the light sheet volume (Figure 4D). Furthermore, the APEX2^+^ cell had dark nuclear and mitochondrial staining with much higher contrast than any of the surrounding cells, confirming we had identified the same APEX2^+^/mCherry^+^ HSPC across multiple imaging modalities. By correlating 3D light sheet and SBEM data, we could confirm lodgement of a single HSPC in the posterior perivascular niche of the KM, allowing us to further define the ultrastructure of this rare cell relative to its surrounding cells in the niche.

To reconstruct the spatial relationships of the single APEX2^+^ HSPC relative to its surrounding niche cells, we performed extensive tracing of cell membranes within the SBEM data using 3D modeling software (IMOD) (Kremer et al., 1996). ECs are generally elongated cells with a large nucleus and little cytoplasm, while MSCs can be distinguish by their granular cytoplasm and a nucleus that occupies three fourths of the cell volume (Tamplin et al., 2015). This morphological analysis revealed the HSPC in the posterior perivascular niche (Figure 5A) is enclosed in a pocket of 5 ECs, and attached to a single MSC, that all directly contact the surface of the HSPC (Figure 5B; Supplementary Video 6). This same configuration of cells (1×HSPC:5xEC:1xMSC) was seen previously in the CHT (Tamplin et al., 2015), demonstrating this cellular structure is conserved between developmental and presumptive adult niches. In addition, a previously unidentified glial-like cell was also in direct contact with the APEX2^+^ HSPC (Figure 5B, C; Supplementary Video 6). Surprisingly, this cell was part of a chain of at least 8 morphologically similar cells that extended throughout the posterior perivascular KM niche. Two putative HSPCs were also present within this niche structure, and together with the APEX2^+^ HSPC, formed a chain of 3 cells in contact (Figure 5B, C). The other 2 HSPCs were ΛPEX2^, suggesting they were not derived by division from the single APEX2^+^ HSPC, and were more likely to be independent HSPC clones that had lodged in the same niche. All 3 HSPCs were in direct contact with the chain of glial-like cells (Figure 5C; Supplementary Video 6). Together, these data show that a single endogenous HSPC in the niche can be in direct physical contact with as many as 8 other cells.

**Figure 5.**
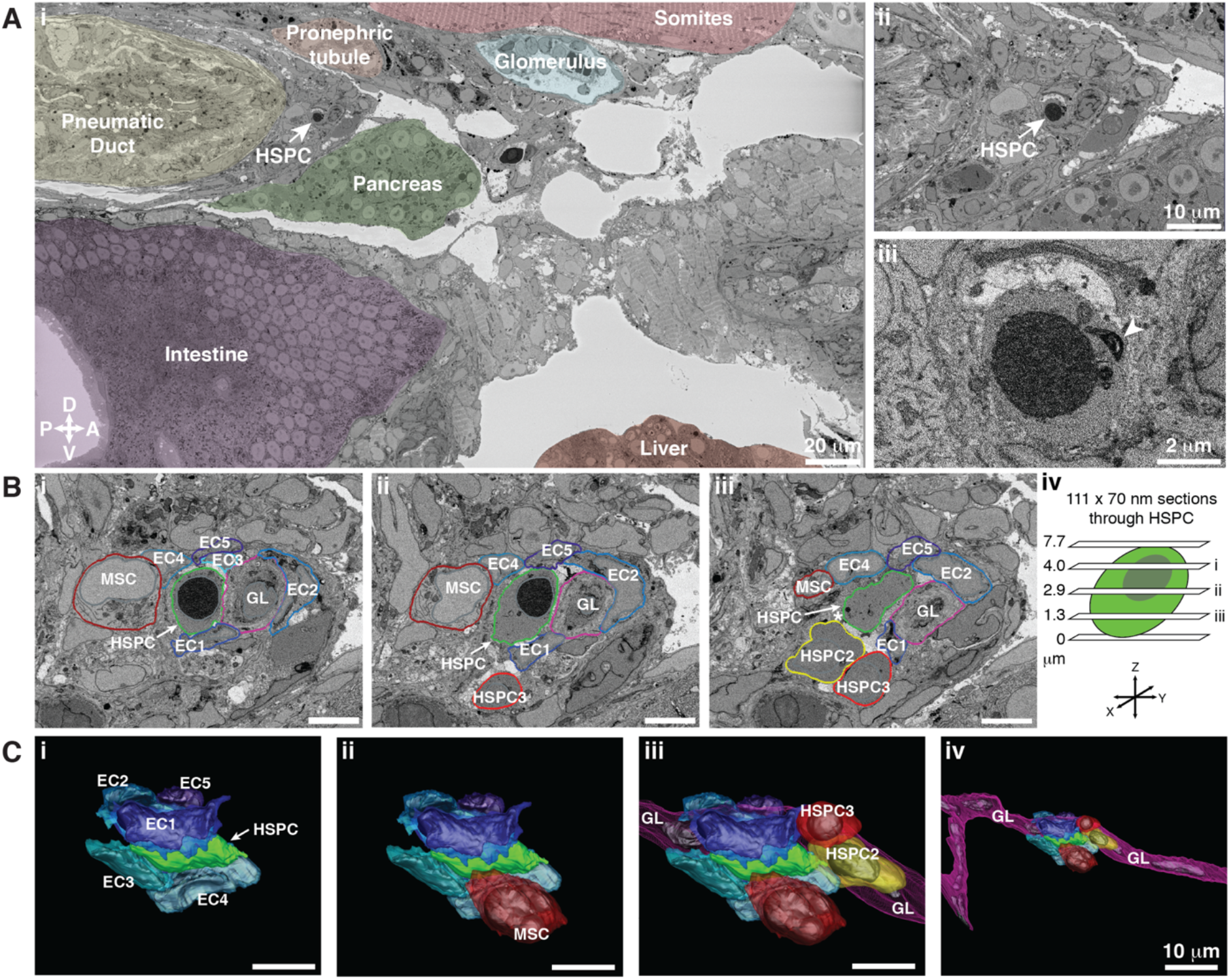
HSPCs lodge in a highly ordered multicellular niche in the perivascular KM. The ultrastructure of a single APEX2^+^ HSPC (white arrow) and its surrounding niche cells are modelled using 3D SBEM. **(A)** The APEX2^+^ HSPC is lodged in the posterior perivascular KM niche. (Ai) Surrounding tissues are labeled, and the HSPC is anterior to the pneumatic duct, dorsal to intestine and pancreas, and ventral to the somites and pronephric tubule. (Dorsal, Ventral, Posterior, Anterior). (Aii) Higher magnification shows the APEX2^+^ HSPC is only 2 cell diameters from the vessel lumen (white area). (Aiii) Full resolution detail of the APEX2^+^ HSPC showing high contrast labeling of the nucleus (APEX2-H2B), mitochondria (mito-APEX2; white arrowhead), and extracellular space dorsal to the cell. **(B)** (i-iii) SBEM sections at different levels through the APEX2^+^ HSPC (white arrows) as shown in the schematic (iv). The HSPC is simultaneously in contact with multiple niche cells: 5 endothelial cells (EC1-5), 1 mesenchymal stromal cell (MSC), and a glial-like (GL) cell. Two unlabeled APEX2-negative putative HSPCs were lodged in the same niche (HSPC2 and HSPC3). HSPC2 is attached to HSPC3, and the APEX2^+^ HSPC (Biii; asterisk). **(C)** 3D rendered models of the APEX2^+^ HSPC (solid green) in contact with niche cells. 3D contours are in the same colors as outlines in (B). The APEX2^+^ HSPC is directly contacted by: (i) 5 ECs; (ii) 1 MSC; (iii) 1 HSPC and a chain of GL-like cells. (iv) The GL-like cell is part of a long continuous chain of similar cells that extends through the niche. Scale bars: 5 μm unless otherwise labeled.

Finally, given this highly ordered HSPC niche structure we observed, we reasoned that an unlabeled HSPC in an independent dataset should be identifiable based on location and cellular morphology alone. We used a second sample that had a single APEX2^+^/mCherry^+^ HSPC that was attached to a vessel wall but was not lodged in the posterior perivascular niche (Figure 6). We searched throughout this posterior perivascular niche region of the 3D SBEM dataset to find unlabeled lodged putative HSPCs. Based on anatomical location and cellular morphology alone, we could construct the same niche model of one putative HSPC in contact with 5 ECs, 1 MSC, and 1 glial-like cell (Figure 7). These data suggest a putative HSPC may be found based on its lodgement in a defined cellular niche structure.

**Figure 6.**
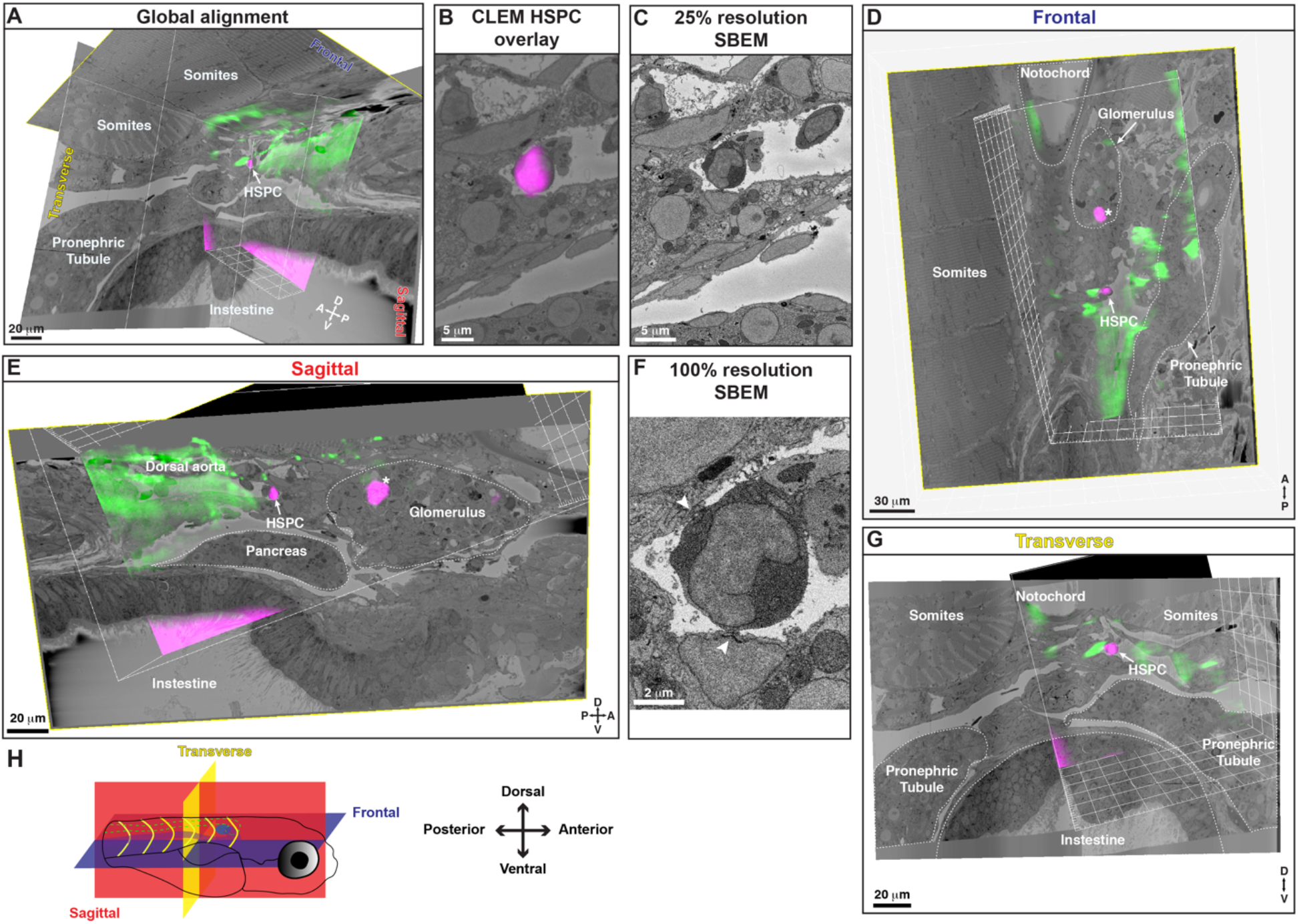
3D alignment of a second dataset also shows localization of a single rare HSPC across multiple imaging modalities. 5 d.p.f. larva. **(A)** Location of the mCherry^+^;APEX2^+^ HSPC (white arrow) in the posterior perivascular niche region, relative to anatomical features such as somites, pronephric tubules, and intestine, aligned in all three orthogonal planes. **(B)** CLEM overlay of mCherry^+^ HSPC imaged with light sheet, overlapping with a single cell in the SBEM data. The target cell is in the lumen of a small vessel in the perivascular niche. Processing and staining of this sample was not optimal, as the nucleus and mitochondria were not resolved in dark contrast on EM sections. **(C)** A single section of SBEM data from the same region shown in (B) at 25% of full resolution. **(D)** Frontal view of the alignment shows notochord, glomerulus, pronephric tubule, somites, mCherry^+^;APEX2^+^ HSPC (white arrow) and a circulating HSPC within the glomerulus (asterix). **(E)** Sagittal view of the alignment shows HSPC (white arrow), circulating HSPC (asterix), glomerulus, dorsal aorta, and pronephric tubule. **(F)** A single section of SBEM data showing the target HSPC at full 100% resolution. The cell is lodged in the lumen of a small vessel and has clear docking sites with adjacent endothelial cells (arrowheads). **(G)** Transverse view shows the HSPC, notochord, pronephric tubules, and intestine. **(H)** Schematic showing the orientation of the larva and the different planes represented: Frontal (blue slice); Sagittal (red slice); Parasagittal (red dotted slice); Transverse (yellow slice).

**Figure 7.**
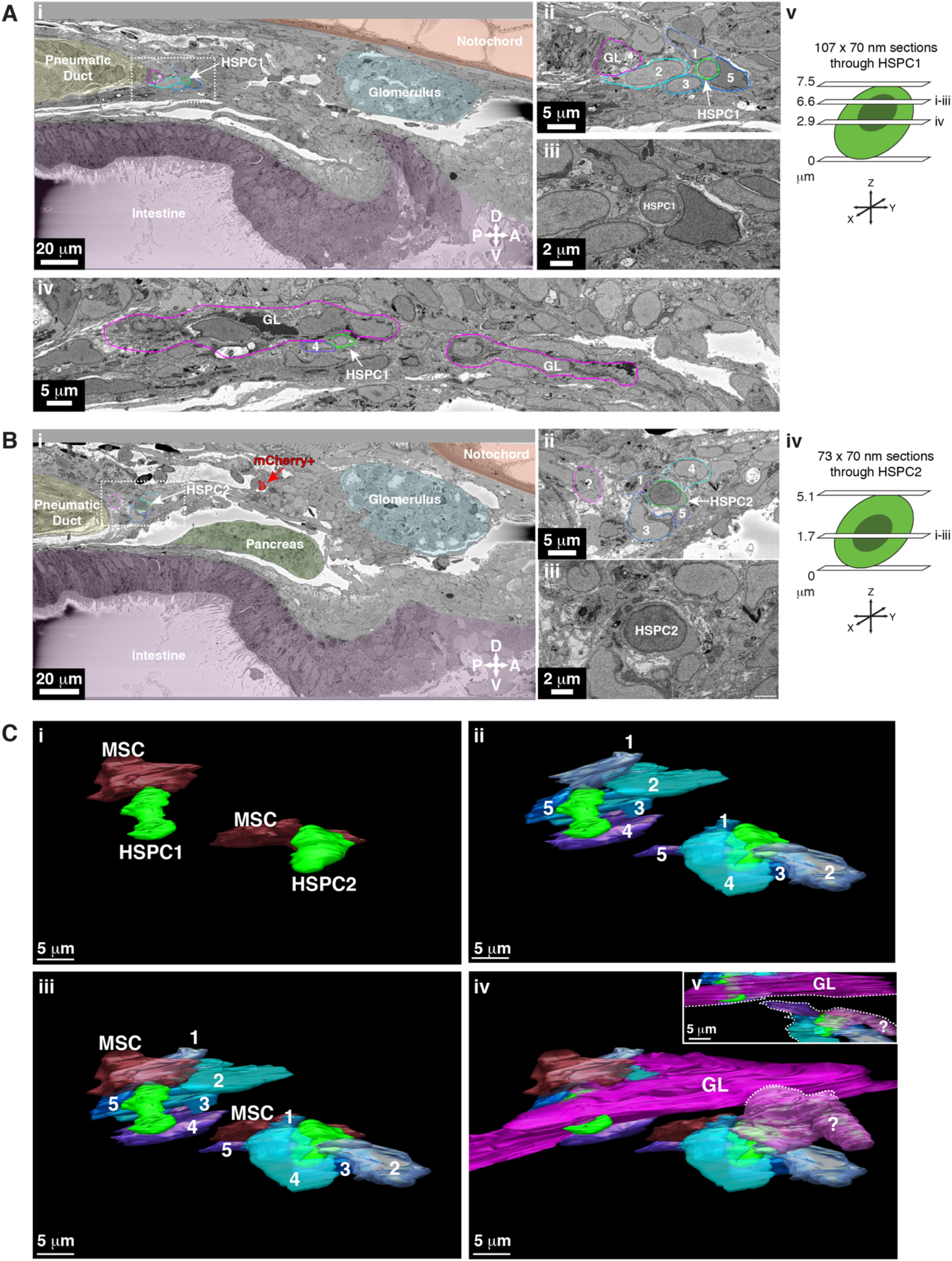
Tracing of unlabeled putative HSPCs in a second SBEM dataset shows conservation of the multicellular niche structure. 5 d.p.f. larva. The ultrastructure of two putative APEX2^-^ HSPCs (based on morphology), and their surrounding niche cells, are modelled using 3D SBEM. The morphological criteria for a putative HSPC were: 1) cell diameter of ~5-7 μm; 2) large round nucleus; 3) scant cytoplasm; 4) ruffled membrane. **(A)** The first putative HSPC (HSPC1; white arrow) we identified was lodged in the posterior perivascular KM niche. (Ai) Surrounding tissues are pseudo-colored. Putative HSPC1 is anterior to the pneumatic duct, dorsal to the intestine, ventral to the notochord, and posterior to the glomerulus. (Aii) Higher magnification of the dotted box shown in (Ai) shows putative HSPC1 is in contact with multiple ECs (1,2,3,5; cell numbers correspond to the traced models in (C)). (Aiii) Full resolution detail of putative HSPC1. (Aiv) The long chain of glial-like (GL) cells (traced in magenta) are in direct contact with putative HSPC1. Cell 4 is one of the ECs shown in the model below (C). (Av) The schematic shows the level of SBEM sections through the putative HSPC. From the bottom of the cell, the section at 6.6 μm is shown in (Ai-iii), and the section at 2.9 μm is shown in (Aiv). **(B)** The second putative HSPC (HSPC2; white arrow) we identified was also lodged in the posterior perivascular KM niche near putative HSPC1. (Bi) Surrounding tissues are pseudo-colored. Putative HSPC2 is anterior to the pneumatic duct, dorsal to the intestine and pancreas, ventral to the notochord, and posterior to the glomerulus. The “mCherry^+^” cell (red with red arrow) was tracked live and aligned using CLEM. We did not see APEX2^+^ high contrast signal in this sample, likely because of sub-optimal DAB staining. (Bii) Higher magnification of the dotted box shown in (Bi) shows putative HSPC2 is in contact with multiple ECs (1,3,4,5; cell numbers correspond to the traced models in (C)). Not shown in this section, the unidentified cell type outlined in light magenta (?) contacts HSPC2 (see Civ). (Biii) Full resolution detail of putative HSPC2. (Biv) The schematic shows the level of the SBEM section through the putative HSPC. From the bottom of the cell, the section at 1.7 μm is shown in (Bi-iii). **(C)** 3D rendered models of putative HSPC1 and HSPC2 (solid green) in direct contact with multiple niche cells. The colors of the 3D contours correspond to the tracing outlines in (A and B). (Ci-iii) Both putative HSPCs are directly contacted by 1xMSC and 5xECs. (Civ) Putative HSPC1 is in contact with a long chain of GL-like cells, like the example shown in Figure 5, which also extends through the niche. Putative HSPC2 is in contact with an unidentified cell type (?). (Cv) The inset shows that the unidentified cell does not connect with the chain of GL-like cells (dotted lines; Civ-v). Note: D=<u>D</u>orsal, V=<u>V</u>entral, P=<u>P</u>osterior, A=<u>A</u>nterior.

To determine the identity of the previously unknown glial-like cells that contacted APEX2^+^ HSPCs, we searched for candidate structures in the kidney region and identified dopamine beta-hydroxylase (dbh) positive ganglia cells to be the most likely candidate (Zhu et al., 2012). Earlier studies in the bone marrow have shown that dopamine signalling is required for normal HSPC trafficking and mobilization (Afan et al., 1997; Katayama et al., 2006; Mendez-Ferrer et al., 2008). In a double transgenic dbh:GFP (Zhu et al., 2012) and Runx:mCherry transgenic line, we observed dbh:GFP^+^ cell projections in contact or close proximity to mCherry^+^ HSPCs (Figure 8A,B; Extended data Figure 9A).

**Figure 8.**
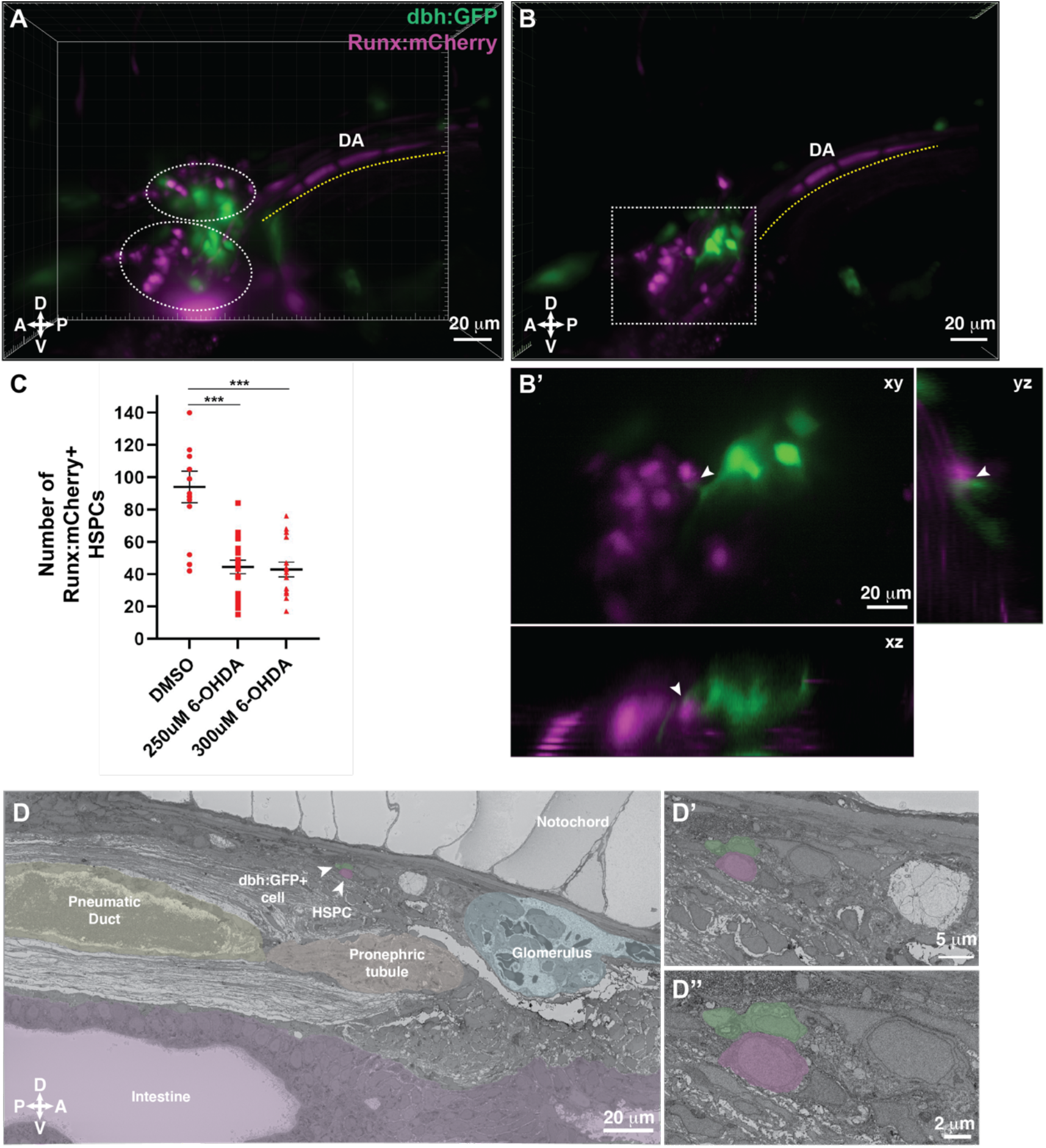
Dopamine beta hydroxylase positive glial-like cells are present within the adult hematopoietic KM niche. **(A)** 3D rendering generated using lightsheet movies of Runx:mCherry;dbh:GFP double transgenic larva shows mCherry^+^ clusters (dotted ovals) in close proximity to GFP^+^ cells. DA, dorsal aorta. (<u>D</u>orsal, <u>V</u>entral, <u>P</u>osterior, <u>A</u>nterior). **(B)** Oblique slice through the 3D volume shows GFP^+^ extensions from the dbh:GFP^+^ cells into the mCherry^+^ HSPC clusters. **(B’)** Detail of the boxed region in B shows contact formation between the GFP^+^ extensions and mCherry^+^ HSPCs (white arrowheads) in all three planes. **(C)** Quantification showing significantly reduced number of Runx:mCherry^+^ HSPCs in 6-OHDA treated transgenic larvae compared to DMSO controls. **(D)** The ultrastructure of a dbh:GFP^+^ cell (labelled green) in proximity of a cell with HSPC-like morphology (labelled magenta) (white arrowheads). Surrounding tissues are labeled, and the dbh:GFP^+^ cell-HSPC pair is posterior to the glomerulus, dorsal to pneumatic duct, pronephric tubule and intestine and ventral to the notochord. **(D’)** Higher magnification shows the dbh:GFP^+^ cell-HSPC pair is only 3 cell diameters from the vessel lumen (white area). **(D”)** Full resolution detail of the dbh:GFP^+^ cell-HSPC pair showing contact formation.

To understand the functional significance of dopamine signalling during KM niche colonization, we treated Runx:mCherry^+^ larvae with 6-hydroxydopamine (6-OHDA), a neurotoxin that induces lesions in dopaminergic neurons (Jackson-Lewis et al., 2012; Matsui and Sugie, 2017; Vijayanathan et al., 2017). We confirmed the efficacy of the drug treatment by reduced locomotor activity (Feng et al., 2014) of the larvae (data not shown). A previous study indicated that disrupting dopamine signalling during early stages of development (5 somite stage till 30 hours post fertilization; h.p.f.) resulted in reduced HSPC numbers within the CHT at 48 h.p.f. (Kwan et al., 2016). Since our aim was to examine the effect of disrupting dopamine signalling on HSPCs during KM niche colonization, we treated larvae with 6-OHDA from 4 to 5 d.p.f. during KM niche colonization. We observed a significant reduction in the number of Runx:mCherry^+^ HSPCs within the niche (Figure 8C; Extended data Figure 9C,D), suggesting a role for dopamine signalling in colonization of the presumptive adult KM niche.

Lastly, we validated the presence of dbh^+^ cells within the KM niche by confocal imaging of dbh:GFP^+^ transgenic larva, followed by SBEM, and then correlation of the two datasets. dbh:GFP^+^ cells aligned with glial-like cells in the KM niche. We identified one glial-like cell in contact with a lodged cell that morphologically resembled an HSPC (Figure 8D; Extended data Figure 9B; Supplementary Videos 7 and 8).

## Discussion

The identification of endogenous stem cells in many tissues remains extremely challenging. We have found it possible to track a single putative HSPC deep in live tissue. Taking advantage of the transparent zebrafish larva and light sheet imaging, we directly observed the kidney marrow niche during the earliest stages of HSPC colonization (4-5 d.p.f.). This is comparable to HSPC seeding of the fetal bone marrow in mammals. Clusters of Runx+ HSPCs were found adjacent to the glomerulus, and single HSPCs were lodged in a perivascular niche. The majority of these perivascular HSPCs were found in pockets of ECs and adjacent or in contact with MSCs, similar to what we previously observed in the CHT (Tamplin et al., 2015). We have also found that ZO-1 positive tight junctions form between HSPCs and surrounding ECs. In *Drosophila* equivalent occluding junctions have been found to regulate niche interactions with hematopoietic cells and germline stem cells (Fairchild et al., 2016; Khadilkar et al., 2017).

To resolve the ultrastructure of HSPCs in the KM niche, we used a genetically encoded fluorescent reporter together with APEX2 that produces a high contrast label in SBEM. This allowed us to match the position of a single labeled cell across multiple imaging modalities. This precise localization of a rare HSPC in a 3D SBEM dataset allowed characterization of its intercellular contacts at the subcellular level. Niche cells immediately surrounding an HSPC formed structures that were highly typical and included five ECs, one MSC, and newly identified glial-like cells. These multicellular structures were distinctive enough that they could be found surrounding unlabeled putative HSPCs. Strikingly, all of these niche cells could simultaneously contact a single HSPC. This highlights the complexity of signals that an HSPC could receive, as well as the challenges ahead to resolve the functional significance of each contact on stem cell regulation.

In the zebrafish larva dopamine beta hydroxylase sympathetic ganglion cells are found in the anterior kidney region at 5 d.p.f. (An et al., 2002; Zhu et al., 2012). We considered that these cells could be the glial-like cells that we found in direct contact with HSPCs. We used additional rounds of CLEM to confirm that dbh:GFP+ cells were adjacent to HSPCs in the perivascular KM niche. Chemical inhibition of dbh during KM colonization significantly reduced the number of Runx+ HSPCs. Our findings are consistent with the established role for neuronal regulation of HSPCs in zebrafish and mammals (Agarwala and Tamplin, 2018). Intriguingly, these dbh+ cells originate from the same neural crest lineage as a subtype of fetal bone marrow MSCs (An et al., 2002; Isern et al., 2014). Together, our results have demonstrated that CLEM is a viable approach to identify single rare stem cells deep in live tissue, and that models built from 3D SBEM data can provide a more complete picture of stem cell-niche interactions.

## Materials and Methods

### Animal care

Adult zebrafish *(Danio rerio)* were maintained at 28C on a 14h:10h light:dark cycle. The Tg(Runx1+23:GFP) (Tamplin et al., 2015) transgenic line used in this study labels single rare HSPCs, and Tg(Runx1+23:mCherry) (Tamplin et al., 2015) and Tg(cd41:GFP) (Lin et al., 2005) label broader HSPC populations. Tg(flk:ZsGreen) (Cross et al., 2003) and Tg(flk:mCherry) (Chi et al., 2008) mark endothelial cells. Tg(cdh17:EGFP) (Zhou et al., 2010) labels the pronephric tubules. Tg(cxcl12/sdf-1a:DsRed2) labels stromal cells (Glass et al., 2011). Tg(dbh:EGFP) (Zhu et al., 2012) marks sympathetic neurons of superior cervical ganglia. Larval zebrafish were raised in petri dishes containing E3 solution (5 mM NaCl, 0.17 mM KCl, 0.33 mM CaCl2 and 0.33 mM MgSO4) until 5 d.p.f. All experiments were performed in accordance with protocols approved by the Institutional Animal Care and Use Committee at the University of Illinois at Chicago.

### Microinjections

Transient transgenesis was established by injecting *draculin:mito-APEX2_p2A_APEX2-H2B_p2A_mCherry_polyA* plasmid together with *tol2* transposase into wild type zebrafish embryos at the single-cell stage. For light sheet imaging, 1 mg ml^-1^ of alpha-bungarotoxin(Swinburne et al., 2015) and 2.5% of Oregon Green dye (Dextran, Oregon Green™ 488; 70,000 MW, Invitrogen) were injected retro-orbitally into the sinus venosus of 5 d.p.f. transgenic larvae to paralyze them and label their vessels, respectively.

### Whole mount immunofluorescence

The 5 d.p.f. Runx1+23:mCherry^+^ zebrafish larvae were fixed overnight in AB buffer containing 4% PFA in 2x fix buffer (234 mM sucrose, 0.3 mM CaCl2 and 1x PBS). Post fixation, the larvae were permeabilized with 0.5% Triton in 1x PBS and washed in deionized water for 2.5 hours. Blocking was performed with 2% BSA in 0.5% Triton/PBS and rabbit polyclonal anti-mCherry primary antibody (Abcam) was used in 1:500 dilution overnight at 4°C. Donkey antirabbit Alexa Fluor 568 (Invitrogen) was used as a secondary antibody at 1:1000 dilution. For phospho-histone H3 (PH3) labeling, mouse monoclonal anti-PH3 (Ser10, clone 3H10) antibody (Millipore Sigma) was used in 1:250 dilution overnight at 4°C. Donkey anti-mouse Alexa Fluor 647 (Invitrogen) was used as a secondary antibody at 1:1000 dilution.

For zonula occludens-1 (ZO-1) antibody staining, the larvae were fixed in 4% PFA overnight followed by 3 PBST (1xPBS with 0.1% Tween 20) rinses. Larvae were dehydrated in increasing methanol/PBST series (25%, 50%, 75%) and stored in 100% methanol at −20°C overnight followed by rehydration in the reversed methanol/PBST series (75%, 50%, 25%). After PBST rinses, they were blocked in 2% serum solution in PBDT (1xPBS with 1% BSA, 1% DMSO and 0.5% Triton) and then incubated in mouse monoclonal ZO1 (clone 1A12) primary antibody (ThermoFisher Scientific) in 1:500 dilution overnight at 4°C. Donkey anti-mouse Alexa Fluor 647 (Invitrogen) was used as a secondary antibody at 1:1000 dilution.

### Drug treatment

The 4 d.p.f. Runx1+23:mCherry^+^ zebrafish larvae were treated with 250 and 300uM of 6-OHDA (Cayman Chemical) for 24 hours. The drug was washed off on 5 d.p.f followed by fixation, tissue clearing and imaging.

### Tissue clearing and confocal imaging of kidney marrow (KM)

To image the KM in fixed samples, following antibody staining, the larvae were embedded in 1% low melting agarose in glass-bottomed dishes (35mm, MatTek Corporation) with the dorsal surface in contact with the glass bottom. To improve visibility through the larvae, optical tissue clearing was performed according to the protocol(Dodt et al., 2007) with some modifications. Embedded samples were washed 6-7 times with 100% methanol followed by overnight incubation at 4°C to dehydrate the tissues. Next, the samples were washed 6-7 times with a 1:1 ratio of 100% methanol and BABB clearing reagent (1part benzyl alcohol: 2 parts benzyl benzoate) and incubated for 30 minutes at room temperature before transferring to 100% BABB. These organic solvents have high refractive indices which result in the solvation of lipids within tissues to align their refractive indices to prevent scattering of light through the tissues (Dodt et al., 2007). Zeiss LSM 880 confocal microscope was used to image the KM. Volume dimensions of 354.25 μm x 354.25 μm x 74 μm was acquired every 2 μm (0.35 x 0.35 x 2 μm/pixel) with a line sequential scan mode and averaging of 2.

For ZO-1 antibody staining, the stained larvae were sectioned in the sagittal plane through vibratome sectioning. Sections of 100 μm were generated that were imaged using Olympus FluoView 1000 at 60x magnification.

### Plasmid construction

The *mito-APEX2_p2A_APEX2-H2B_p2A_mCherry* cassette was synthesized on a kanamycin resistant pUC19 backbone (GenScript) and assembled using Multisite Gateway Cloning (Invitrogen). The draculin 5’ entry promoter construct (Mosimann et al., 2015) was assembled with middle entry vector *pME_mito-APEX2_p2A_APEX2-H2B_p2A_mCherry,* 3’ entry vector Tol2kit #302*p3E-polyA* and destination vector Tol2kit #394*pDestTol2pA2* (Kwan et al., 2007).

### Light sheet Imaging

For light sheet imaging, stage-matched transgenic zebrafish larvae were paralyzed by retro-orbital injection of alpha-bungarotoxin (Swinburne et al., 2015) and embedded in 1% low melting point agarose within thin capillary tubes. The KM was illuminated with a light sheet from one axis, and fluorescence was detected at a perpendicular axis. Larvae were imaged with a 20x objective using the Zeiss Light sheet Z.1 for Single Plane Illumination Microscopy (SPIM). The capillary tube was inserted into the sample chamber to release the larva into the chamber containing E3 while remaining attached to the capillary tube through the agarose layer. The larva was rotated such that the lateral surface of the larva closer to the edge of the agarose layer was perpendicular to the detection objective to optimize fluorescence detection. The laser path was aligned to illuminate the KM from one axis. Z-stacks were acquired through the entire KM with 1 μm spacing in less than 1 minute and time-lapse videos were recorded to track the circulating HSPCs entering the KM. Through a full 360° rotation of the larva relative to the detection lens, only about 5° were optimal for light emission from the sample and image acquisition of the KM.

### Image analysis

Image processing and rendering were done using Imaris.v.9.1 (Bitplane), IMOD(Kremer et al., 1996), and ImageJ/Fiji (Schindelin et al., 2012). Imaris was used to count the number of cells within the KM niche. Imaris “Spot” module was used first to specify the region of interest around the KM in confocal images followed by automated counting of the spots generated corresponding to the single HSPCs. The accuracy of the spots generated was further confirmed by using the manual mode of “Spot” function. “Co-localization” feature on Imaris was used to identify PH3/mCherry double positive cells. Distances between different cell pairs were measured using Imaris. Imaris “Add image to” function was used to align 3D rendered model generated from light sheet data over 3D rendered SBEM dataset using anatomical markers as reference. Orthogonal views through the two datasets in the XY, YZ and XZ planes were used for the alignment. ImageJ/Fiji was used to orient and define the region of interest within the microCT stack. For tracking and aligning ROIs across imaging modalities, we used an in-house landmark-based software that facilitates the 3D registration between volumes with an iterative procedure that progressively establishes sets of common markers. Once the relative transformations are determined, a composite volume containing all the image modalities can be visualized as a single Tiff stack with Fiji (Imaris, Amira or any other rendering software). In the perfectly aligned high-resolution SBEM dataset, IMOD was used to manually trace individual cells of interest. Separate objects were defined by drawing contours around the plasma membrane of the target cell as it moved slice by slice in the Z plane. Individual contours were meshed with *imodmesh* to reveal 3D reconstructions of the cells of interest. In this way, HSPCs, ECs, MSCs, GLs and their nuclei were identified and represented in the 3D model. Fluorescent image levels and background were adjusted, using the default “background subtraction” (Bitplane Imaris), threshold adjustment, and/or brightness/contrast adjustment (Fiji/ImageJ, Bitplane Imaris, Zeiss Zen, Adobe Photoshop). SBEM brightness/contrast adjustment was performed using IMOD and/or Fiji, Imaris, Adobe Photoshop.

### *draculin:mito-APEX2_p2A_APEX2-H2B_p2A_mCherry^+^* zebrafish sample preparation

5 d.p.f. zebrafish larvae were prepared for MicroCT and SBEM as previously described (Deerinck et al., 2010). Briefly, immediately after light sheet imaging, larvae were fixed in 2.5% glutaraldehyde and 4% paraformaldehyde in 0.15 M cacodylate buffer (CB, pH 7.4) at 4°C overnight. After removing the fixative, larvae were treated for 15 minutes with 20 mM glycine in 0.15M CB, on ice to quench the unreacted glutaraldehyde. Preincubation was done in 2.5 mM DAB solution (25.24 mM stock in 0.1 M HCl) in 0.15 M CB for 1 hour on ice. For staining, 0.03% H_2_O_2_ containing DAB solution was added to the larvae on ice. The reaction was monitored every 5-10 minutes and stopped when the desired intensity was achieved. The staining buffer was washed off with 5x washes using 0.15 M CB. Then, larvae were washed with 0.15 M CB and then placed into 2% OsO4/1.5% potassium ferrocyanide in 0.15 M CB containing 2 mM CaCl2. The larvae were left for 30 min on ice and then 30 min at room temperature (RT). After thorough washing in double distilled water (ddH_2_O), larvae were placed into 0.05% thiocarbohydrazide for 30 min. Larvae were again washed and then stained with 2% aqueous OsO4 for 30 min. Larvae were washed and then placed into 2% aqueous uranyl acetate overnight at 4°C. Larvae were washed with ddH_2_O at RT and then stained with 0.05% en bloc lead aspartate for 30 min at 60°C. Larvae were washed with ddH_2_O and then dehydrated on ice in 50%, 70%, 90%, 100%, 100% ethanol solutions for 10 min at each step. Larvae were then washed twice with dry acetone and placed into 50:50 Durcupan ACM:acetone overnight. Larvae were transferred to 100% Durcupan resin overnight. Larvae were then flat embedded between glass slides coated with mould-release compound and left in an oven at 60°C for 72 h.

### dbh:GFP^+^ zebrafish sample preparation

To facilitate the 3D correlation between the different imaging modalities, DRAQ5, a DNA binding fluorescent dye was used. Briefly, the 5dpf dbh:GFP^+^ transgenic zebrafish larvae were fixed with 0.5% glutaraldehyde and 2% paraformaldehyde in 0.15M cacodylate buffer (CB, pH 7.4) for 2.5 hours and 100 μm thick sagittal sections were collected and incubated in DRAQ5 (1:1000, Cell Signaling Technology) on ice for an hour. Confocal images of GFP and DRAQ5 signals were collected on a Leica SPE II confocal microscope with a 20X oil-immersion objective lens using 488 nm and 633 nm excitation. Following, the samples were preincubated in 2.5 mM DAB solution (25.24 mM stock in 0.1 M HCl) in 0.15 M CB for 30 min on ice. Illumination of DRAQ5 locally induces the generation of reactive oxygen, that subsequently triggers the diaminobenzidine (DAB) polymerization by oxidation resulting in darkening of the nuclei (Ou et al., 2017). Therefore, the photo-oxidation of DAB by DRAQ5 was done in 2.5 mM DAB in 0.15 M CB using a solar simulator (Spectra-Physics 92191–1000 solar simulator with 1600 W mercury arc lamp and two Spectra-Physics SP66239–3767 dichroic mirrors to remove infrared and ultraviolet wavelengths), while bubbling oxygen in the solution. The light was filtered through 10cm square bandpass filters (Chroma Technology Corp.) for illumination at 615 nm (40 nm band pass). The reaction was monitored every 20 minutes and stopped when the desired darkening in the nuclei was achieved. Larvae were then washed 5 times with 0.15 M CB and incubated in 2% OsO4/1.5% potassium ferrocyanide in 0.15 M CB containing 2 mM CaCl2 to get an EM visible stain. The larvae were left for 30 min on ice and then 30 min at room temperature (RT). After thorough washing in double distilled water (ddH_2_O), larvae were placed into 0.05% thiocarbohydrazide for 30 min. Larvae were again washed and then stained with 2% aqueous OsO4 for 30 min. Larvae were washed and then placed into 2% aqueous uranyl acetate overnight at 4°C. Larvae were washed with ddH_2_O at RT and then stained with 0.05% en-bloc lead aspartate for 30 min at 60°C. Larvae were washed with ddH_2_O and then dehydrated on ice in 50%, 70%, 90%, 100%, 100% ethanol solutions for 10 min at each step. Larvae were then washed twice with dry acetone and placed into 50:50 Durcupan ACM:acetone overnight. Larvae were transferred to 100% Durcupan resin overnight. Larvae were then flat embedded between glass slides coated with mould-release compound and left in an oven at 60°C for 72 h.

### MicroCT and Serial Block-face scanning Electron Microscopy (SBEM)

The MicroCT tilt series were collected using a Zeiss Xradia 510 Versa (Zeiss X-Ray Microscopy) operated at 80 kV (88 μA current) with a x20 magnification and 0.872 μm pixel size. MicroCT volumes were generated from a tilt series of 2401 projections using XMReconstructor (Xradia). SBEM were accomplished using Merlin SEM (Zeiss, Oberkochen, Germany) equipped with a Gatan 3View system and a focal nitrogen gas injection setup (Deerinck et al., 2018). This system allowed the application of nitrogen gas precisely over the block-face of ROI during imaging with high vacuum to maximize the SEM image resolution. Even a minimal accumulation of charge on the specimen surface resulted in poor image quality and distortions in the non-conductive biological samples. This could happen through 3,000 sequential images of the charge-prone zebrafish block-face due to image jitter. Deerinck et al. recently introduced a new approach to SBEM that uses focal nitrogen gas-injection over the sample block surface to eliminate the charging, while allowing the high vacuum to be maintained in the specimen chamber, resulting in a tremendous improvement in image resolution, even with specimens that were not intensely heavy-metal stained (Deerinck et al., 2018). Images were acquired in 3 kV accelerating voltage and 0.5 μsec dwell time; Z step size was 70 nm; raster size was 30k x 18k and Z dimension was ~ 3,000 image samples. Volumes were collected using 40% nitrogen gas injection to samples under high vacuum. Once volumes were collected, the histograms for the slices throughout the volume stack were normalized to correct for drift in image intensity during acquisition. Digital micrograph files (.dm4) were normalized and then converted to MRC format. The stacks were converted to eight bit and volumes were manually traced for reconstruction using IMOD (Kremer et al., 1996).

## Data availability

SBEM datasets generated in this study will be deposited in a publicly accessible data archive. Newly generated plasmids will be deposited in Addgene.

## Acknowledgements

This work was supported by grants from the NIH NHLBI (R01HL142998), NIDDK (K01DK103908), American Society of Hematology (Junior Faculty Scholar Award; O.J.T.), and the American Heart Association (Grant #19POST34380221; S.A.). Light sheet imaging was performed at the Integrated Light Microscopy Core Facility at the University of Chicago with the help of Christine Labno and Vytas Bindokas. The authors wish to thank: Dr. Daniela Boassa for her assistance in EM probe design with APEX2; Tom Deerinck, Steven Peltier, and Tristan Shone for technical advice with Focal CC; Iain A. Drummond for feature identification in SBEM sections.

## Author contributions

S.A. and O.J.T. performed zebrafish experiments. K.Y.K. and E.A.B. performed microCT and SBEM experiments. S.A., K.Y.K., S.P., S.J., Y.E.K, I.A.D, and O.J.T. analysed data. S.A., O.J.T, K.Y.K., E.A.B., and M.H.E. designed experiments. M.H.E. and O.J.T supervised the project.

## Materials & Correspondence

Correspondence and requests for materials should be addressed to O.J.T.

---

**Extended Data Figure 1.**
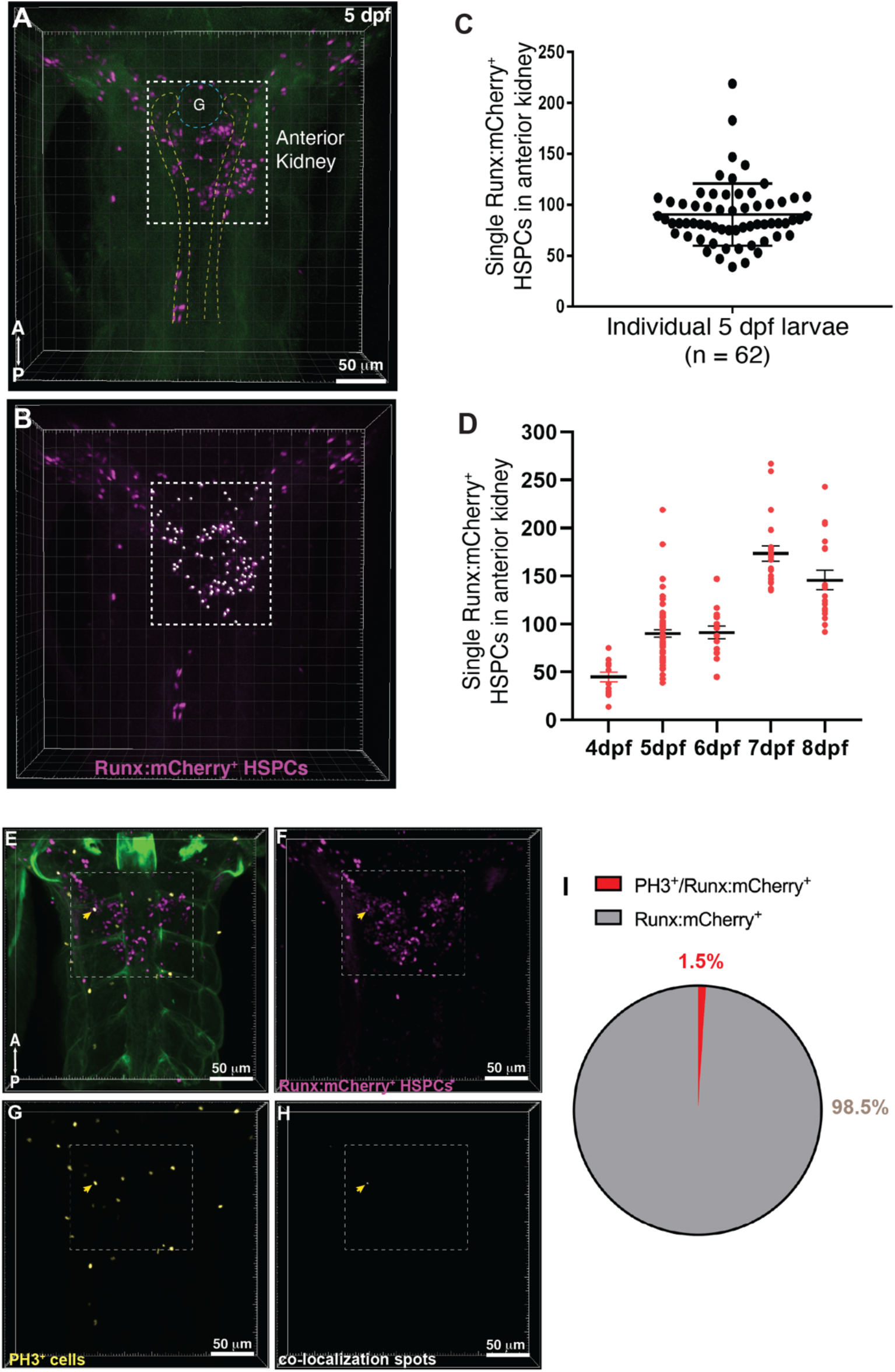
**(A)** 3D rendering of a confocal stack shows the dorsal view of a fixed 5 d.p.f. Runx:mCherry (magenta) transgenic zebrafish larva imaged after optical tissue clearing. Runx:mCherry^+^ HSPC clusters are located in the anterior kidney region (dotted box; G, glomerulus, dotted blue circle; pronephric tubules, dotted yellow lines). Green signal is autofluorescence. **(B)** Quantification of Runx:mCherry^+^ HSPCs (white spots in dotted box). **(C)** Quantification shows a mean of ~100 HSPCs at 5 d.p.f.. **(D)** Number of Runx:mCherry^+^ HSPCs within the niche increases from 4d.p.f. and peaks at 7d.p.f. followed by a reduction at 8d.p.f.. (4 d.p.f., n = 13; 5 d.p.f., n = 62; 6 d.p.f., n = 15; 7 d.p.f., n =21; 8 d.p.f., n = 18). **(E-I)** 3D rendering of confocal stack shows the dorsal view of a fixed 5 d.p.f. Runx:mCherry **(E)** transgenic zebrafish larva showing Runx:mCherry^+^ HSPCs **(F)** stained with PH3 antibody **(G)** to label the proliferating cells. **(H)** mCherry and PH3 staining co-localize in one HSPC within the niche as seen by the co-localization spot (yellow arrowheads). **(I)** Pie chart for PH3^+^ and Runx:mCherry^+^ double positive cells amongst Runx:mCherry^+^ cells shows that less than 2% of HSPCs proliferate at this stage (n = 11).

**Extended Data Figure 2.**
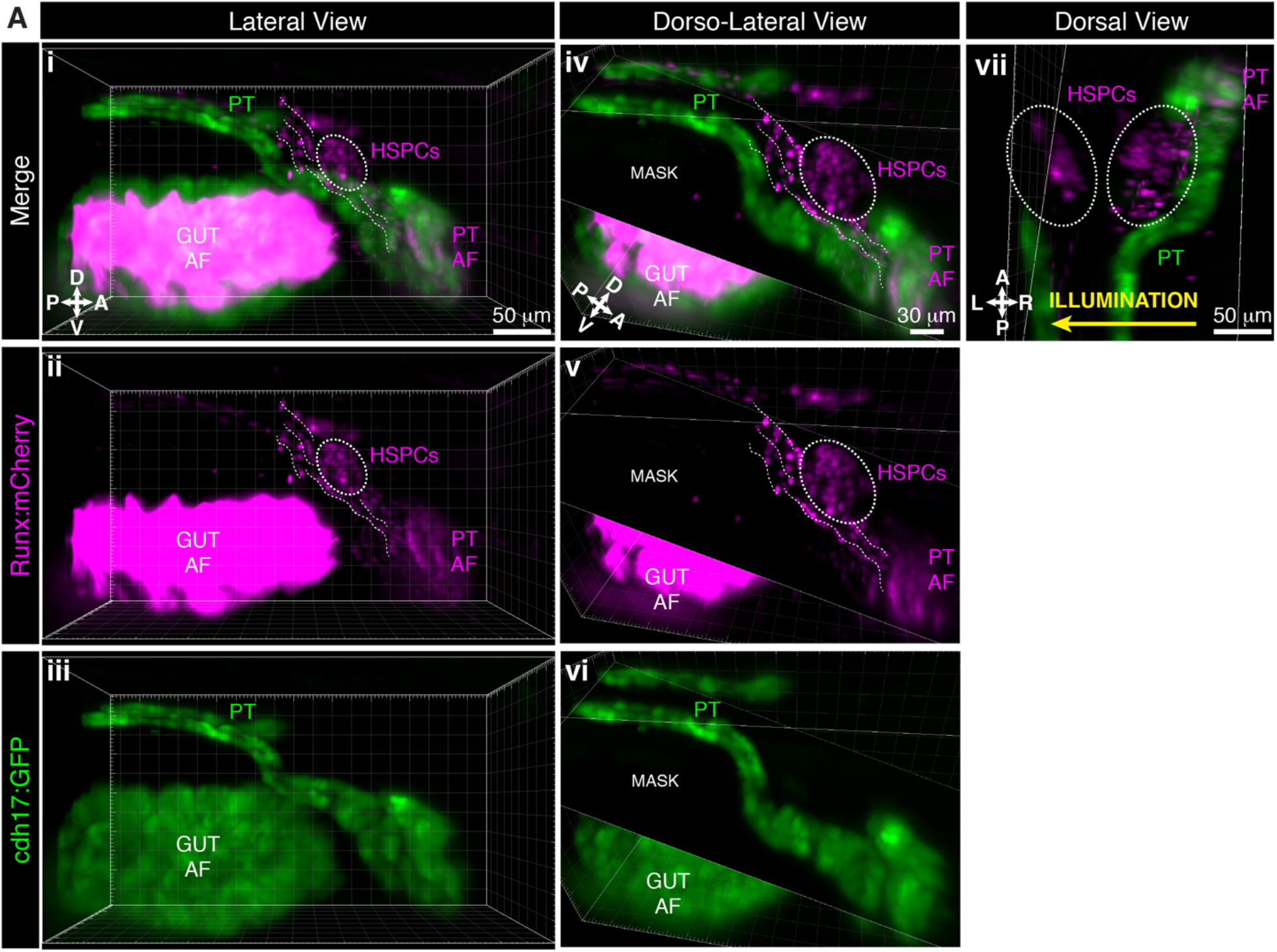
Light sheet live imaging shows HSPC clusters are mediolateral to the proximal pronephric tubules (PT). Larva is 5 d.p.f. (Ai-Aiii) Lateral and (Aiv-Avi) dorsolateral views of Runx:mCherry and cdh17:GFP double transgenics labeling the HSPCs and PT, respectively. HSPC cluster (dotted oval) relative to proximal PT. The dotted lines show cells in circulation. ‘Mask’ is an oblique slice on Imaris to block the background autofluorescence (AF) from the gut. (Avii) Dorsal view of the same larva shows two HSPC clusters located mediolaterally to proximal PT. Light sheet imaging illuminated this sample from the right side, showing how signal and resolution is poorer further from the light source.

**Extended Data Figure 3.**
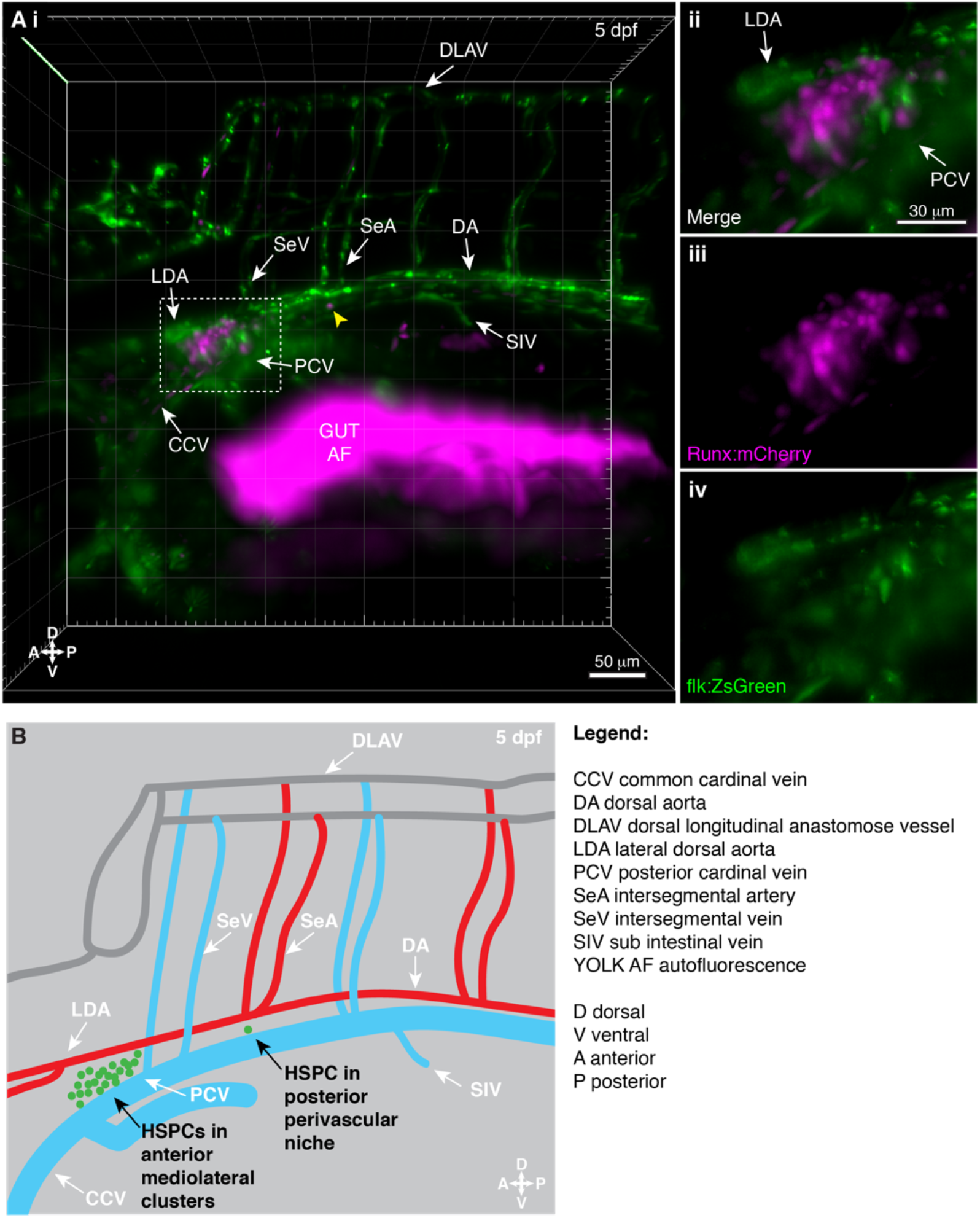
Light sheet live imaging of the anterior kidney reveals HSPCs in distinct perivascular regions. Larva is 5 d.p.f.. **(A)** (i) 3D projection of the anterior kidney in Runx:mCherry^+^ (HSPCs) and flk:ZsGreen^+^ (ECs) double transgenic larva imaged using light sheet microscope. HSPC cluster (mCherry^+^) is located between ZsGreen^+^ LDA and PCV (white dashed box). Single mCherry^+^ HSPC is in a perivascular niche that is posterior to the HSPC cluster (yellow arrowhead). (Aii) Detail of the mCherry^+^ HSPC cluster located between ZsGreen^+^ LDA and PCV (merged), (Aiii) Runx:mCherry^+^ HSPCs alone, and (Aiv) flk:ZsGreen^+^ LDA and PCV alone. **(B)** Schematic showing the location of HSPCs (green dots) relative to the vasculature in the KM. HSPCs form anterior mediolateral clusters or are located as single cells within the posterior perivascular niche.

**Extended Data Figure 4.**
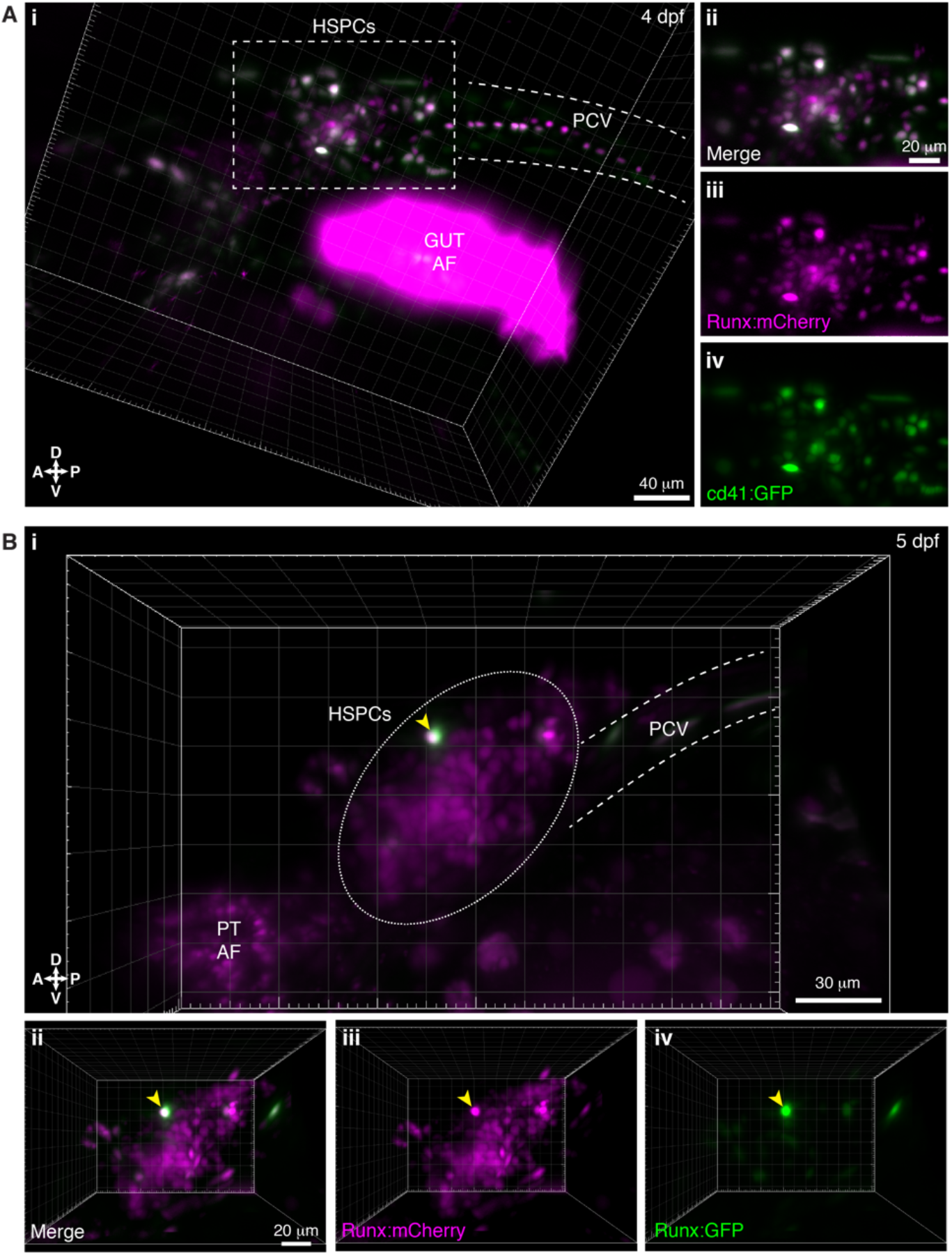
Light sheet live imaging of overlap of HSPC transgenic reporter lines. **(A)** Runx:mCherry and cd41:GFP double positive larva (4 d.p.f.). (i) 3D projection shows overlap between the Runx:mCherry and cd41:GFP positive populations (white dashed rectangle). (ii-iv) Detail of the HSPC cluster. **(B)** Runx:mCherry and Runx:GFP double positive larva (5 d.p.f.). (i) 3D projection shows overlap between the Runx:mCherry and Runx:GFP positive populations (yellow arrowhead), with Runx:mCherry more broadly expressed in HSPCs, as previously published(Tamplin et al., 2015). (ii-iv) Detail of the HSPC cluster. PT: pronephric tubules, AF: autofluorescence, PCV: posterior cardinal vein.

**Extended Data Figure 5.**
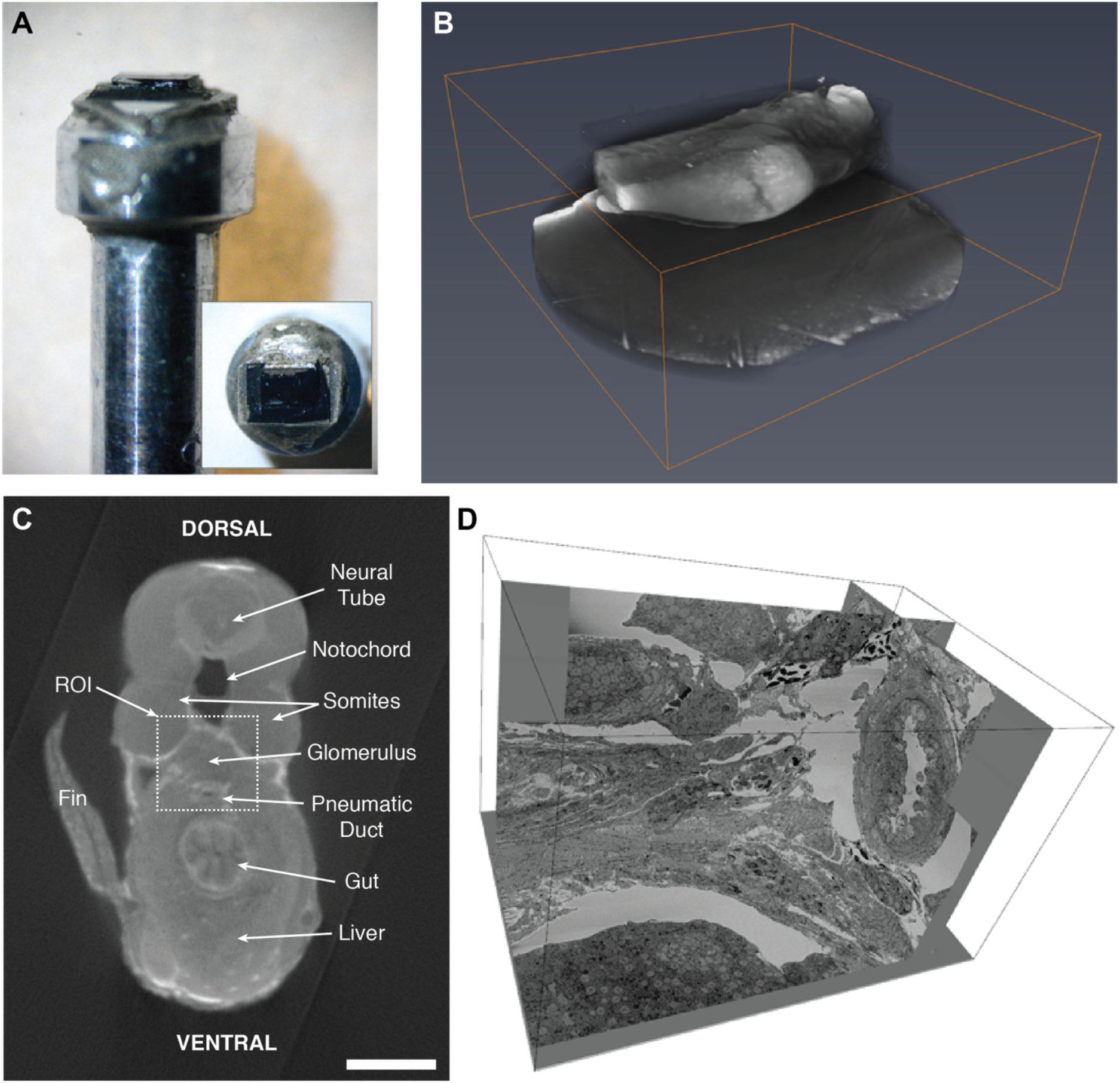
Method for mounting, aligning and trimming the sample for SBEM. **(A)** The zebrafish tissue block on the aluminium pin is completely opaque to light following staining for SBEM. **(B)** MicroCT reveals details within the SBEM-stained tissue and helps to find the exact region of interest (ROI). **(C)** Single section of microCT data set labeled with anatomical features. Scale bar: 100 μm. **(D)** SBEM volume (dimensions: 31×22×30 μm) and a reconstruction of HSPCs and multicellular niche.

**Extended Data Figure 6.**
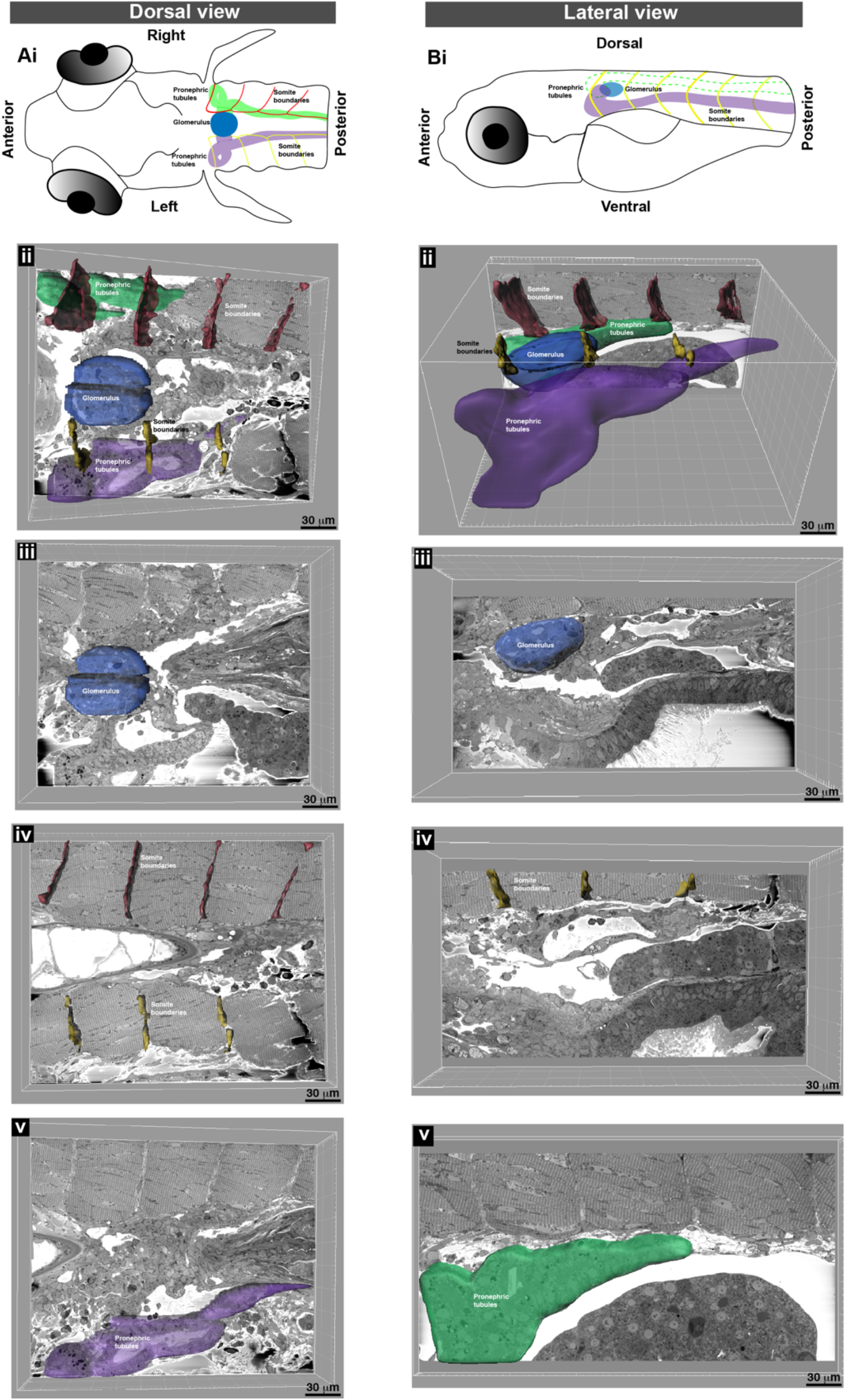
Segmentation and 3D surface rendering identifies anatomical features within the SBEM datasets. **(A)** (i) Schematic of the dorsal view of zebrafish larvae. (ii) Oblique slice through 3D SBEM data shows the right and left pronephric tubules, glomerulus and the right and left somite boundaries. Different frontal planes through the SBEM showing the (iii) glomerulus, (iv) somite boundaries, and (v) left pronephric tubules. **(B)** (i) Schematic of the lateral view of zebrafish larvae and (ii) Imaris oblique slice through SBEM data shows pronephric tubules, glomerulus and somite boundaries. Different sagittal planes through the SBEM showing the (iii) glomerulus, (iv) left somite boundaries and (v) left pronephric tubule. Visualization performed using Imaris software (Bitplane).

**Extended Data Figure 7.**
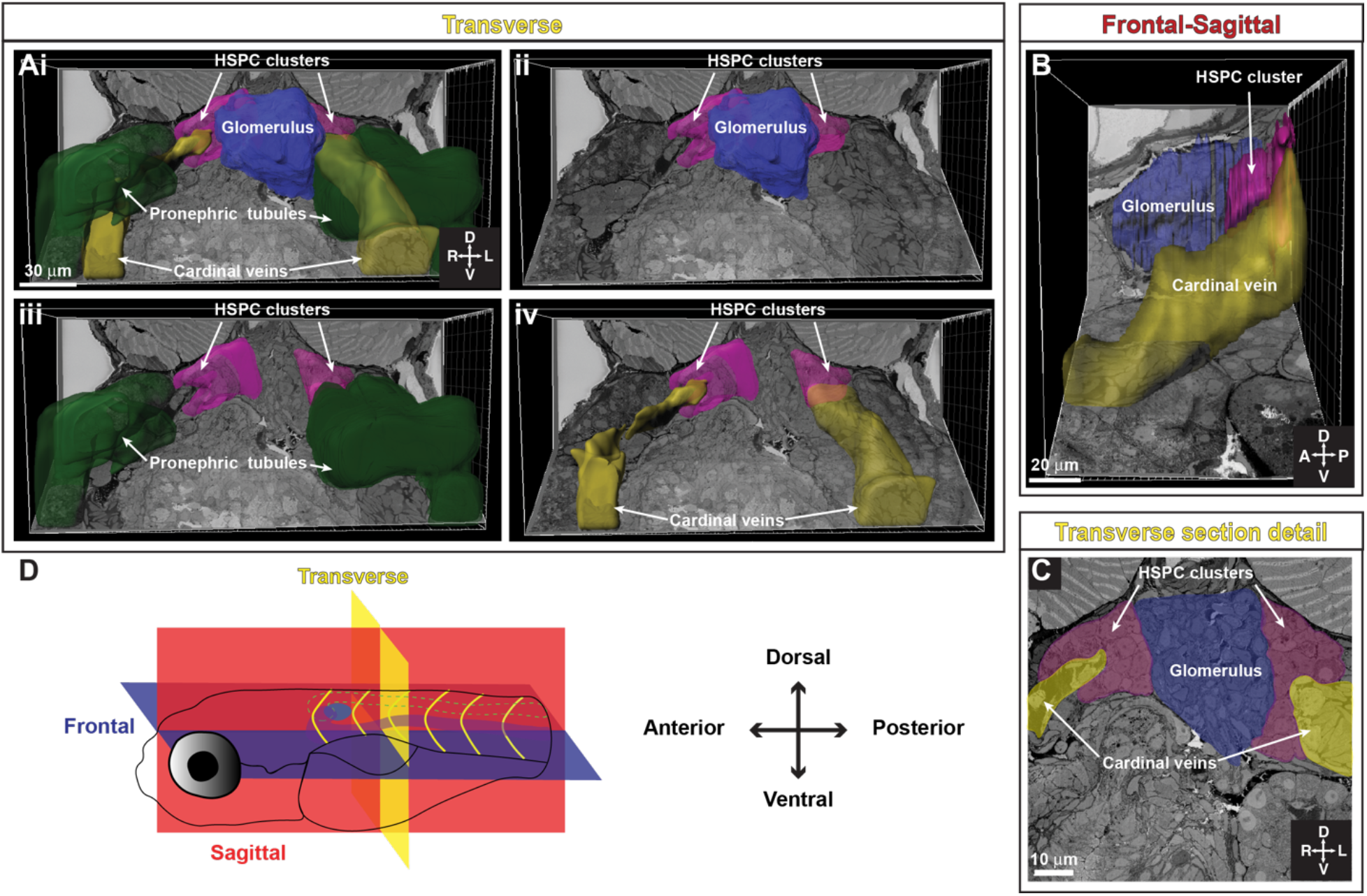
Tracing of SBEM shows location of anterior HSPC clusters relative to the glomerulus, pronephric tubules, and cardinal veins. **(A)** (i) Oblique slice along the transverse view of 3D SBEM of zebrafish larvae shows the location of anterior HSPC clusters relative to the pronephric tubules, cardinal veins, and the glomerulus. Anterior HSPC clusters are located on either side of the glomerulus (ii), medio-lateral to pronephric tubules (iii), and the cardinal veins (iv). **(B)** Oblique slice shows the frontal-sagittal view of 3D SBEM of zebrafish larvae where the HSPC cluster is located between the glomerulus and the cardinal vein. **(C)** Detail of a single section in the transverse view shows the location of bilateral HSPC clusters, located between the glomerulus and the cardinal veins. **(D)** Schematic showing the orientation of the larva and the different planes represented.

**Extended Data Figure 8.**
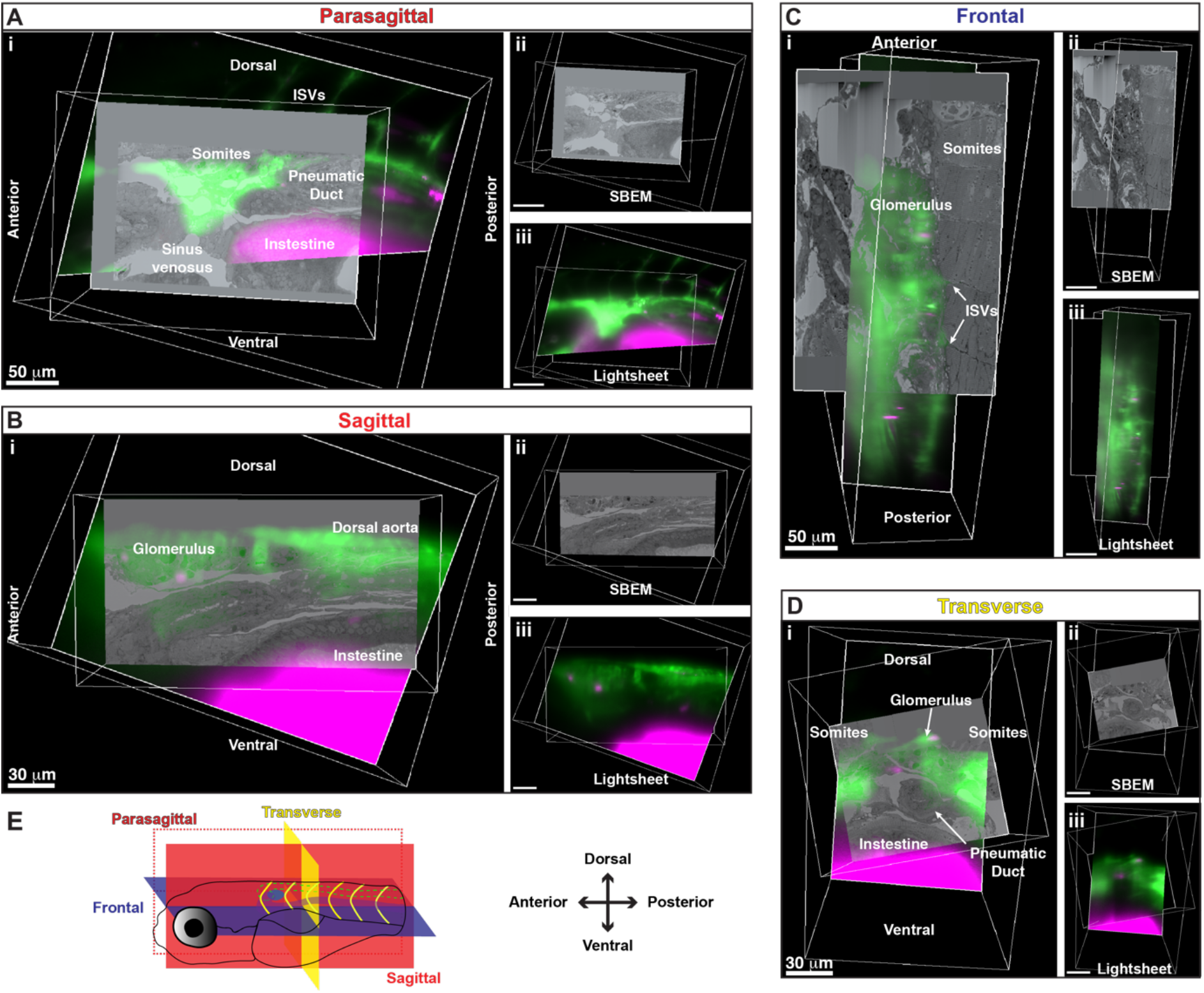
Alignment of 3D rendered models from SBEM and light sheet live imaging shows anatomical features in all three planes. **(A)** Parasagittal view shows somites, pneumatic duct, intestine, sinus venosus and inter-segmental vessels (ISVs). **(B)** Sagittal view shows glomerulus, dorsal aorta and intestines. **(C)** Frontal view shows the glomerulus, ISVs, and somites. **(D)** Transverse view shows the glomerulus, somites, pneumatic duct, and intestine. **(E)** Schematic showing the orientation of the larva and the different planes represented. (Ai, Bi, Ci, Di) Merged 3D models of light sheet and SBEM data. (Aii, Bii, Cii, Dii) 3D SBEM volume alone. (Aiii, Biii, Ciii, Diii) 3D light sheet volume alone.

**Extended Data Figure 9.**
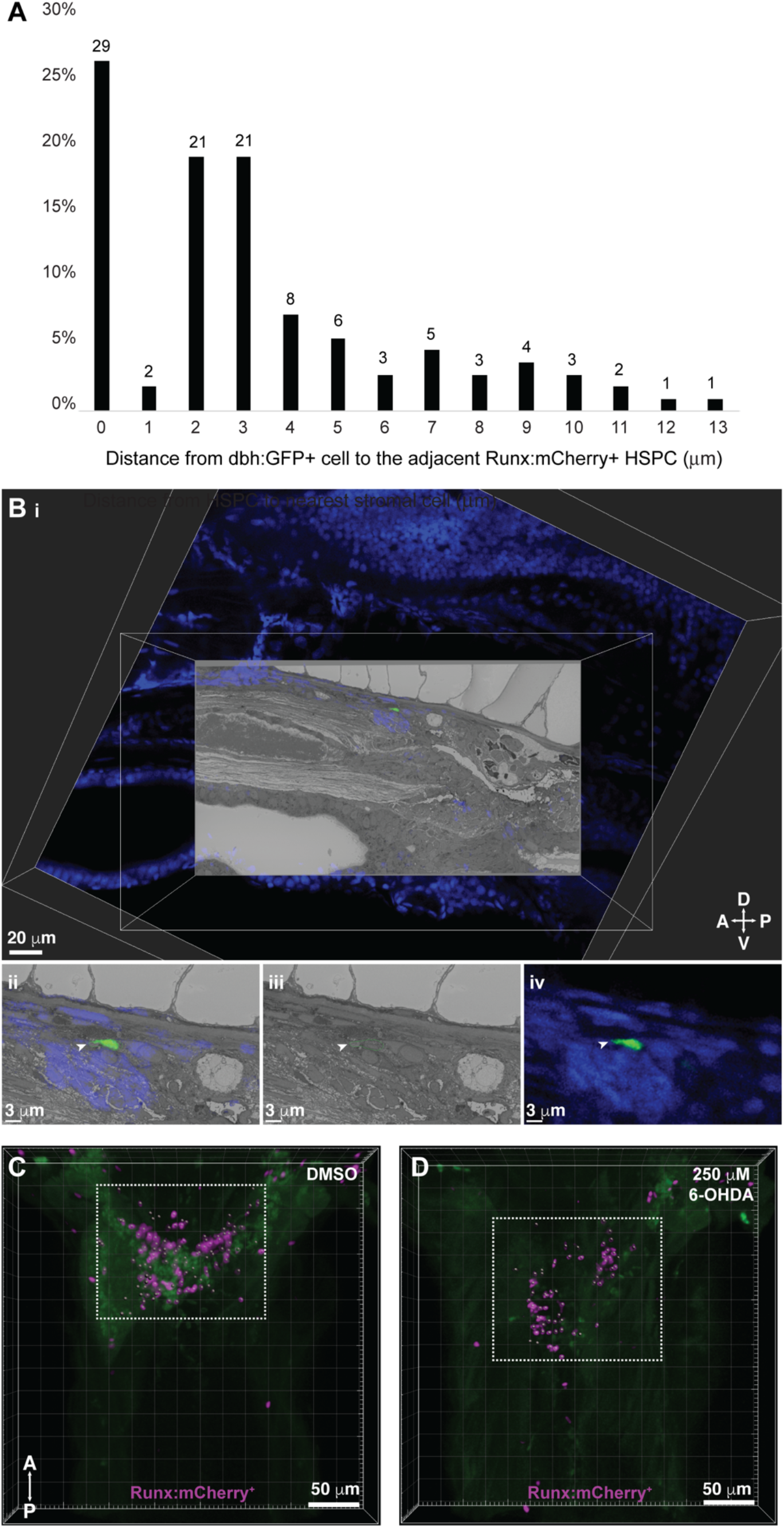
CLEM shows that dbh:GFP^+^ cells are present within the KM and neurotoxin targeting dopaminergic neurons reduces HSPC colonization of the KM niche. **(A)** Quantification of distances measured between dbh:GFP^+^ cells and Runx:mCherry^+^ HSPCs shows that almost 30% of HSPCs are in contact with dbh:GFP^+^ cells and the remaining are within 13 μm. Numbers above the columns indicate the cell numbers counted in each group (n=10). **(B)** 3D alignment of confocal and SBEM datasets localizes dbh:GFP^+^ cell to the KM niche. (i) Orthogonal view shows the alignment between confocal and SBEM datasets in XY plane. White arrowhead points to the single dbh:GFP^+^ cell in the aligned confocal and SBEM datasets (ii), green outlined dbh:GFP^+^ cell in SBEM data (iii), and dbh:GFP^+^ cell in confocal data. **(C,D)** 3D rendered confocal stacks show reduced number of Imaris software-defined “spots” within the anterior kidney region (in the white dotted box) corresponding to reduced Runx:mCherry^+^ HSPCs upon 250 μM 6-OHDA treatment (D), as compared to DMSO treated controls (C). Green is autofluorescence.

## Supplementary Videos

**Video 1. Depth-coded projection of a time-lapse video shows migrating mCherry^+^ HSPCs occupying the KM niche.** Runx:mCherry^+^ zebrafish larva embedded in low melting agarose was aligned to the laser path from the left side to illuminate the KM, which results in a distinctly visible mediolateral HSPC cluster on the left compared to the right side. Depth-coding is along a rainbow spectrum, with red-orange-yellow on the left side, green-blue in the midline, and violet on the right side. Circulating HSPCs are seen entering the anterior mediolateral HSPC clusters. Larva is 105 hours post fertilization (h.p.f.) at the start of imaging. Frames were captured every 2 minutes. The video was rendered at 2 frames per second (f.p.s.). Timestamp is hours:minutes after start of imaging. SeA, intersegmental artery; SeV, intersegmental vein; DA, dorsal aorta; Gut AF, autofluorescence; D, dorsal; V, ventral; A, anterior; P, posterior.

**Video 2. Single circulating HSPCs interact with the posterior perivascular niche region.** Time-lapse video shows Runx:GFP^+^ HSPCs (green) in circulation through flk:mCherry^+^ vessels (magenta). Occasionally, HSPCs slow and lodge in the posterior perivascular niche region. Larva is 5 d.p.f. at the start of imaging. Frames were captured every 12.5 seconds. The video was rendered at 15 frames per second (f.p.s.). Timestamp is minutes:seconds after start of imaging. LDA, lateral dorsal aorta; SeA, intersegmental artery; SeV, intersegmental vein; DA, dorsal aorta; PCV, posterior cardinal vein; D, dorsal; V, ventral; A, anterior; P, posterior.

**Video 3. Transiently labeled HSPCs lodge within the perivascular niche.** Time-lapse movie shows a single lodged mCherry^+^ HSPC (magenta; yellow arrowhead) in a transient draculin:2APEX2_mCherry^+^ injected larva. The cell is lodged ventral to the DA and anterior to the PD. Dextran-conjugated Oregon green dye is injected to label the vessels (green). Larva is 5 d.p.f. at the start of imaging. Frames were captured every 2 minutes. The video was rendered at 10 frames per second (f.p.s.). ISV, intersegmental vessels; DA, dorsal aorta; PD, pneumatic duct; Gut AF, autofluorescence.

**Video 4. microCT stack acquired through a fixed zebrafish larva.** microCT is an intermediate step between light sheet imaging and SBEM that is used to orient the sample and define our region of interest to proceed with SBEM.

**Video 5. All sections (>3,000) of the SBEM data set shown in Figures 3 and 4.** Rendered at 60 frames per second (f.p.s.). 1/10 full resolution.

**Video 6. HSPCs lodge in a multicellular niche structure in the perivascular KM.** Orthogonal views through the SBEM data in all three planes shows the dense KM region. APEX2^+^ staining of HSPCs allows us to distinguish these cells from the other cell types. IMOD software was used to trace the outlines of the HSPC and the surrounding niche cells such as endothelial cells, mesenchymal stromal cells and glial-like cells that form an HSPC niche unit.

**Video 7. Correlation of confocal and microCT stacks.** Orthogonal views through the correlated stack in XY plane. Magnification 20X.

**Video 8. Dbh:GFP^+^ cells form a part of the KM niche.** Orthogonal views through correlated confocal and SBEM data in the XY plane shows that dbh:GFP^+^ cells are located within the KM niche. Rendered at 10 frames per second (f.p.s.). 1/10 full resolution of SBEM.

